# Circuit mechanisms underlying chromatic encoding in *Drosophila* photoreceptors

**DOI:** 10.1101/790295

**Authors:** Sarah L. Heath, Matthias P. Christenson, Elie Oriol, Maia Saavedra-Weisenhaus, Jessica R. Kohn, Rudy Behnia

**Author notes:** contributed equally (Sarah L. Heath, Matthias P. Christenson).

## Abstract

Spectral information is commonly processed in the brain through generation of antagonistic responses to different wavelengths. In many species, these color opponent signals arise as early as photoreceptor terminals. Here, we measure the spectral tuning of photoreceptors in *Drosophila*. In addition to a previously described pathway comparing wavelengths at each point in space, we find a horizontal-cell-mediated pathway similar to that found in mammals. This pathway enables additional spectral comparisons through lateral inhibition, expanding the range of chromatic encoding in the fly. Together, these two pathways enable optimal decorrelation of photoreceptor signals. A biologically constrained model accounts for our findings and predicts a spatio-chromatic receptive field for fly photoreceptor outputs, with a color opponent center and broadband surround. This dual mechanism combines motifs of both an insect-specific visual circuit and an evolutionarily convergent circuit architecture, endowing flies with the unique ability to extract chromatic information at distinct spatial resolutions.

## Introduction

Color vision is an important source of visual information, enhancing our recognition of objects in complex visual fields. How is wavelength information extracted by the brain? A single type of photoreceptor cannot distinguish wavelength independently of the intensity of light because different spectral distributions of varying intensity can give rise to the same photoreceptor output (Rushton, 1972). Color percepts can only be extracted by comparing the output from at least two photoreceptors with different spectral sensitivities. This comparison is apparent in color opponent neurons, which receive antagonistic inputs from different photoreceptor types and therefore exhibit opposing responses to different ranges of wavelengths (Shevell and Martin, 2017). Our understanding of the neural processes that lead to our perception of colors therefore critically depends on our understanding of color opponent signals and the underlying circuits that establish them.

Much of what we know about the properties of color opponent neurons comes from work done in the tri-chromatic retina of primates. There, the signals from L, M and S cones, named for their sensitivity in the long, middle and short wavelength regions of the spectrum, are combined by two main types of opponent retinal ganglion cells (RGCs): the so-called “red-green” neurons, which compare the activity of M and L photoreceptors, and “blue-yellow” neurons, which compare the activity of S and L+M photoreceptors (reviewed in Conway et al. 2010). Because cone photoreceptors are arranged in a 2D lattice, lateral interactions are essential for establishing these opponent signals in the retina. This results in spectrally opponent signals in RGCs which compare chromatic information between neighboring points in visual space through center-surround interactions. Interestingly, the two axes of opponency - “red-green” and “blue-yellow” - encoded at the level of RGCs have been shown to correspond to an optimal decomposition of S, M and L cone sensitivities (Buchsbaum and Gottschalk, 1983). This allows the retina to remove the correlations introduced by the high degree of overlap between cone sensitivities and more efficiently transmit spectral information to downstream visual circuits.

Opponent signals have been measured across the animal kingdom, reinforcing the importance of this operation in color circuits across evolution. *Drosophila melanogaster* has emerged as a genetically tractable system to study circuit level mechanisms of color vision (Gao et al., 2008; Karuppudurai et al., 2014; Melnattur et al., 2014; Schnaitmann et al., 2013, 2018; Yamaguchi et al., 2010), and color opponent signals have been measured at the axonal terminals of cone-like photoreceptors in the fly brain (Schnaitmann et al., 2018). However, unlike the 2D lattice photoreceptor arrangement found in mammals, the light sensing rhabdomeres of fly cone-like photoreceptors R7 and R8 are positioned one on top of each other (Hardie, 1985) (Figure 1A). This architecture allows photoreceptors in each optical unit, or ommatidium, to absorb photons emanating from the same point in visual space. A specialized circuit taking advantage of this configuration was recently described to generate color opponent signals through reciprocal inhibition exclusively between pairs of R7 and R8 photoreceptors from a single ommatidium (Schnaitmann et al., 2018), allowing for pixel-by-pixel comparison of wavelengths. Because of the spectral composition of the fly eye, these intra-ommatidial interactions impose specific constraints on the types of spectral comparisons that the circuit can make. Indeed, there are two types of ommatidia in the main part of the fruit fly eye, that are distributed in a stochastic pattern (65% “yellow”, 35% “pale”, Figure 1A, D) (reviewed in Behnia and Desplan 2015). “Pale” ommatidia express the short-UV-sensitive Rh3 rhodopsin in R7 and the blue-sensitive Rh5 in R8. “Yellow” ommatidia express the long-UV-sensitive Rh4 rhodopsin in R7 and the green-sensitive Rh6 in R8. An opponent mechanism purely based on intra-ommatidial interactions therefore defines two separate color opponent channels, both comparing spectral information along a UV vs visible axis.

**Figure 1.**
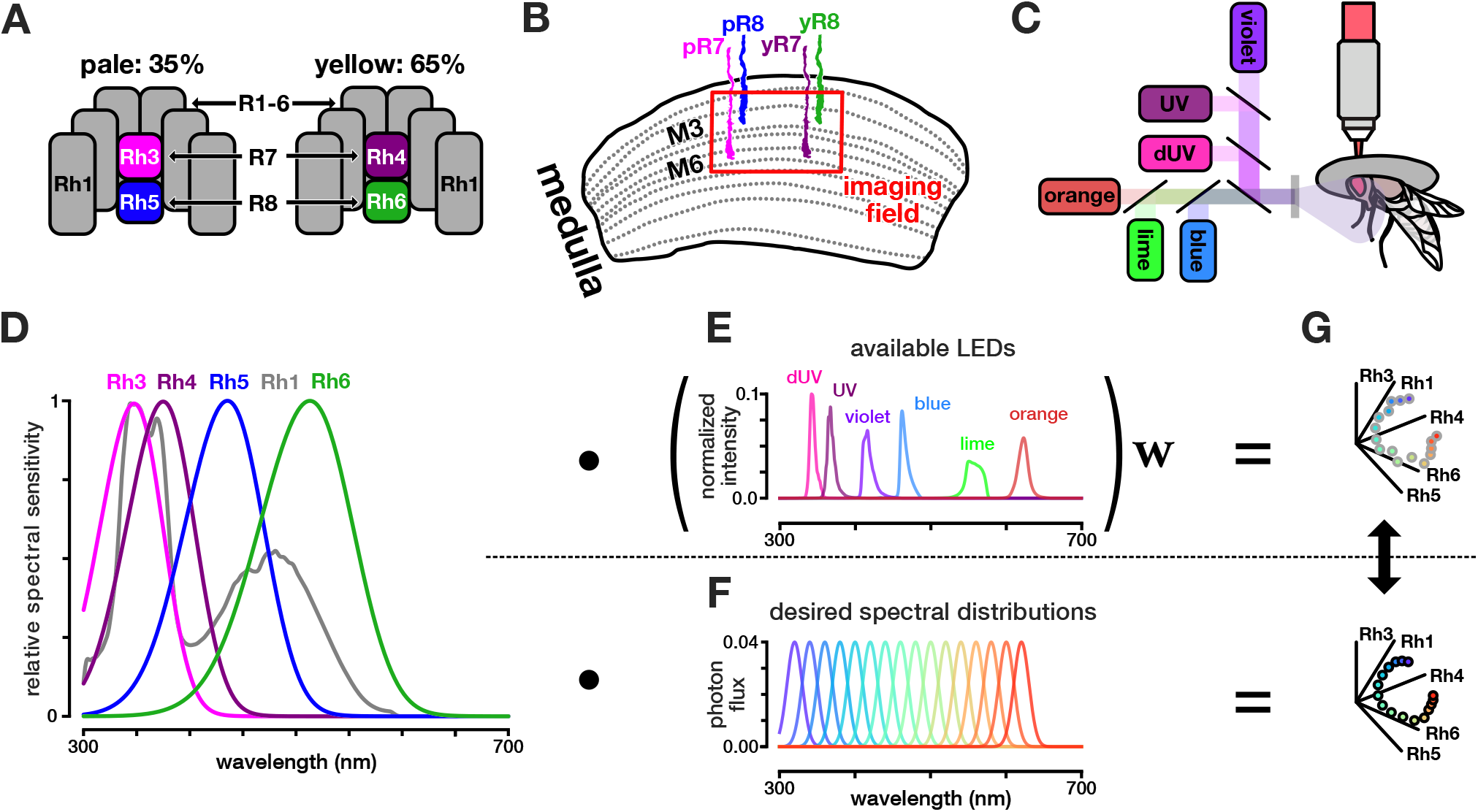
Experimental setup and stimulus design. **A.** Spectral composition of pale and yellow ommatidia of the *Drosophila* eye. Pale ommatidia express Rh3 and Rh5 in R7 and R8, respectively. Yellow ommatidia express Rh4 and Rh6 in R7 and R8, respectively. R1-6 all express Rh1. **B.** Photoreceptors in *Drosophila* project from the retina into the optic lobe. Our imaging experiments target the axon terminals of R7 and R8 in the medulla at the level of layers M6 and M3, respectively. **C.** Two-photon imaging set up. The fly is secured facing a screen, and LED sources are combined using a custom color mixer to form a single collimated beam. **D.** Relative spectral sensitivity of opsins expressed in the fruit fly retina (adapted from Salcedo et al. 1999). **E.** Normalized photon flux across the wavelength spectrum, corresponding to the various LEDs used for stimuli. **F.** Desired set of spectral distributions to test to build a spectral tuning curve. **G.** For any given single wavelength in **F**, we calculate the relative photon capture (*q*) for all five opsins by integrating over the opsin sensitivities in **D**, and plot a vector in photon capture space. We then simulate the single wavelength with combinations of the available LEDs in **E** that most closely recreate that vector (see methods for details).

This architecture has the advantage of allowing chromatic information to be extracted at the full resolution of the eye, similarly to achromatic pathways driven by R1-6 photoreceptors (Clandinin and Zipursky, 2000), which express the broadband opsin Rh1 (Figure 1A, D). However, it does not allow for additional comparisons to be made in the spectral domain, such as those between the blue and green part of the spectrum, which appear to be used behaviorally (Melnattur et al., 2014; Schnaitmann et al., 2013; Yamaguchi et al., 2010), and which may be beneficial in terms of efficient signal processing. Lateral interactions between R7s and R8s from neighboring ommatidia, akin to those mediated by horizontal cells in the mammalian retina (Thoreson and Mangel, 2012), would allow for increased resolution of chromatic pathways in the spectral domain and provide the fly with more flexible mechanisms for encoding chromatic information.

Here, we measure the spectral tuning of all four types of wavelength-specific photoreceptors in the fly visual system. We find that each R7 and R8 photoreceptor type displays specific and distinct wavelength opponent properties, which cannot be explained solely by previously described reciprocal inhibition within single ommatidia. At the circuit level, we show that indirect antagonistic interactions between R7s and R8s from neighboring ommatidia also contribute to shaping the spectral tuning of all photoreceptor outputs. Furthermore, we find that these indirect interactions are mediated by the horizontal-cell-like medulla interneuron Dm9. These indirect interactions enable additional comparisons in the spectral domain which correspond to optimal decorrelation of the spectral sensitivities of *Drosophila* opsins. In addition, we show that photoreceptor inputs are integrated linearly, which allows us to build a linear recurrent model constrained by the underlying circuit interactions: reciprocal inhibitory interactions between R7/R8 in the same ommatidium, R7/R8 inhibitory inputs onto Dm9, and excitatory feedback from Dm9 onto all R7s/R8s. This model accurately predicts our observed responses, while also showing that electron-microscopy-based synaptic count provides an accurate proxy for synaptic weight in this early processing step in color circuits. Our circuit model predicts a receptive field for R7 and R8 outputs with a broad-band surround superimposed on a color-opponent center, combining the motifs of both an evolutionarily convergent circuit architecture and an insect-specific visual circuit.

## Results

### R7 and R8 rhabdomeric responses are transformed into opponent outputs through interactions between photoreceptor types

Color opponent responses are established via antagonistic interactions of inputs from different types of photoreceptors. In the case of *Drosophila* R7 and R8, rhabdomeric responses of these photoreceptors in the eye can be considered inputs, and their color opponent axonal responses in the medulla can be considered outputs (Figure 2A). To understand how inputs are combined to give rise to color opponent outputs, our first goal was to measure and compare the rhabdomeric and the axonal spectral tuning properties of these photoreceptors.

**Figure 2.**
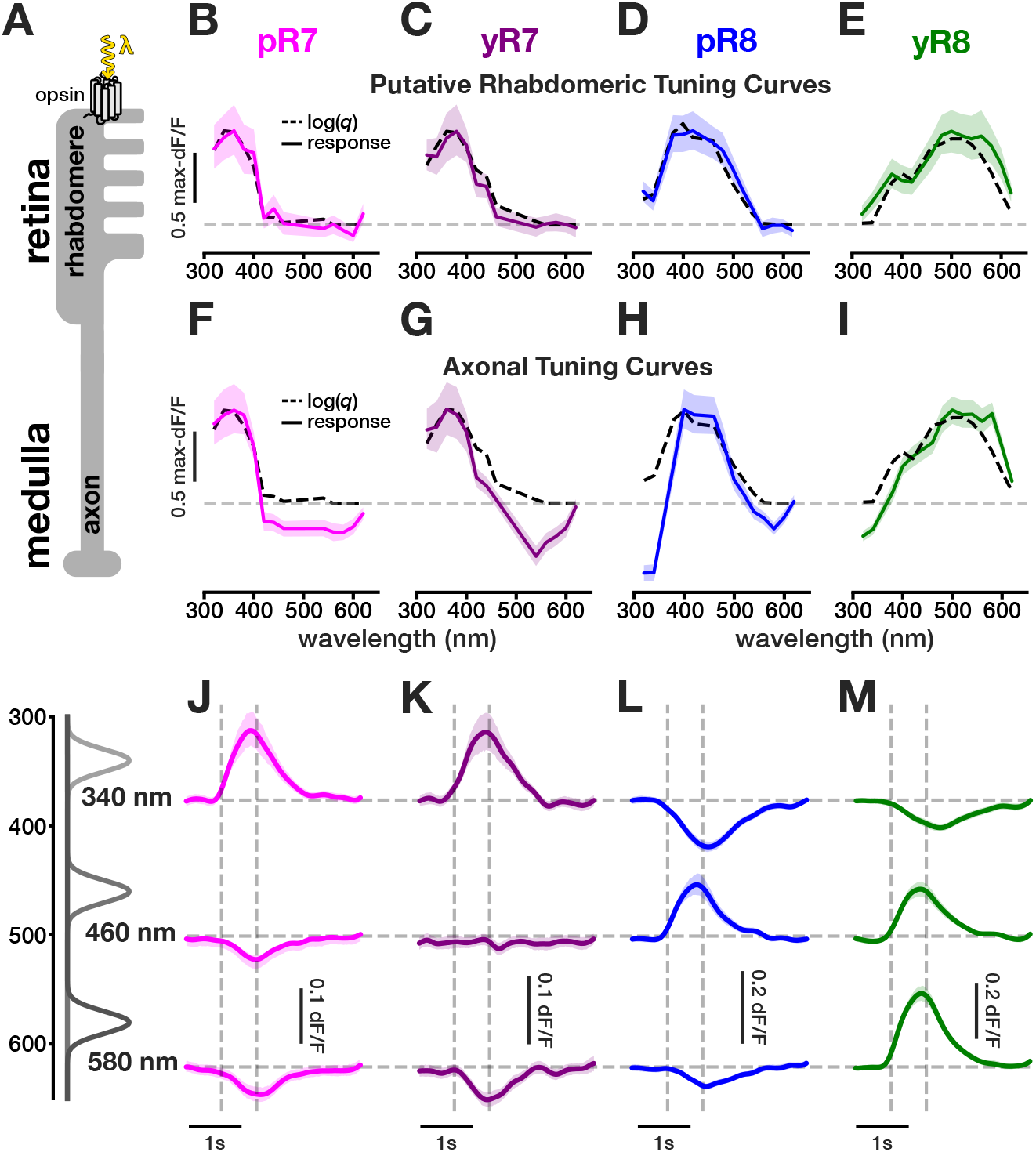
R7 and R8 rhabdomeric responses are transformed into opponent outputs. **A.** In *Drosophila* photoreceptors, light (*λ*) is absorbed in the retina by rhodopsin molecules at the level of the rhabdomeres, where phototransduction takes place. Photoreceptors project their axons to the medulla where synaptic interactions occur. **B-E.** NorpA, an essential component of the phototransduction cascade, was restored in *norpa-* blind flies in individual photoreceptor types. This allowed for measurement of putative rhabdomeric spectral tuning in photoreceptor axons by eliminating interactions from other cell types. Maxnormalized responses of R7/R8 axons were measured across simulated wavelengths to construct spectral tuning curves. Ns= 106 ROIs (8 flies), 96(8), 69(7), and 26(4), respectively. Dashed black lines represent the log (*q*). Colored lines represent the mean photoreceptor response. Shaded region represents the 95% confidence interval. Dashed grey lines represent baseline fluorescence. **F-I.** Max-normalized spectral tuning curves constructed using the amplitudes of measured responses of R7 and R8 axons in wild type flies. Ns= 152(8), 134(6), 138(7), and 129(6), respectively. **J-M.** Average GCaMP6f responses of R7 and R8 axons in wild type flies to 0.5 second flashes of three simulated wavelengths. Vertical dashed grey lines represent onset and offset of light presentation.

*In vivo* two-photon imaging of genetically targeted GCaMP6f in R7 and R8 photoreceptors allows for straightforward measurement of their axonal outputs in the M6 and M3 layers of the medulla, respectively (Figure 1B). However, we could not visualize rhabdomeres in the eye with our imaging setup. Instead, we used genetic tools to make indirect measurements of rhabdomeric responses. Because these responses are transformed into axonal outputs through interactions with other photoreceptor types (Schnaitmann et al., 2018), we reasoned that measurements at the axonal level in mutant flies where these interactions are abolished correspond to putative pure rhabdomeric responses. For this set of experiments, we isolated these responses in mutant flies where only the imaged photoreceptor type is active, effectively preventing external inhibitory input from other photoreceptor types. This is done by functionally rescuing phototransduction in single photoreceptor types in the blind *norpa-* mutant background by driving expression of UAS-NorpA with Rhodopsin-Gal4 drivers (Wernet et al., 2012).

In order to compare rhabdomeric and axonal tuning, we developed a method to measure spectral tuning curves (Figure 1D-G). Specifically, we measured neuronal responses to a range of relatively narrow-band light sources of equal photon flux (photons per second per unit area; *E* =photon flux per mole) spanning the fly’s visible spectrum. Instead of using a system with a large number of different light sources, we devised a method that allows us to measure tuning curves using only a limited number of LEDs. For a given light source, each photoreceptor type will “capture” a specific number of photons. This number, or photon capture, is calculated as a function of each opsins sensitivity and the spectrum of the light source (Kelber et al. 2003; Renoult et al. 2017; see methods; Equation 5). We simulated the effect of this particular light source on the fly eye by showing a combination of the six LEDs in our stimulus set-up. By using least-squares regression to calculate the appropriate intensities for each LED (Equation 7), we evoked the same capture in each of the five photoreceptor types (y/pR7, ypR8 and R1-6) as the intended narrow-band light source (see methods and Figure S1 for details on implementation and accuracy). Measuring responses to these simulated light sources across the spectrum allowed us to construct spectral tuning curves for a given cell type. All experiments were performed in light adapted conditions where the simulated light source is presented over a background light. Because we anticipated measuring opponent waveforms at the level of photoreceptor outputs, we used single wavelength dominant backgrounds (UV for R7s and blue for R8s), which have the advantage of highlighting opponent signals. In these conditions, GCaMP6f fluorescence was indeed increased at baseline, allowing decreases to be readily measured.

As expected from the spectral sensitivity of the opsins they express, the putative rhab-domeric responses we measured show UV sensitivity in p/yR7 peaking at 360 nm and 380 nm, respectively, blue sensitivity in pR8 peaking at 420 nm, and blue/green sensitivity in yR8 peaking at 500 nm (Figure 2B-E). These neural responses are directly related to spectral sensitivities of the opsins that these photoreceptors express. It was previously shown that a logarithmic transformation of photon capture corresponds to the transformation of light absorption of a photoreceptor by the phototransduction cascade (Henderson et al., 2000; Juusola and Hardie, 2001). We thus compared the tuning curves we obtained to the log of the relative photon capture log(*q*) in each rhodopsin, specifically calculated for the presented stimuli. We found that the measured tuning curves closely match the calculated log(*q*). This result shows that log(*q*) is a reliable estimate of rhabdomeric responses in this system, which we will subsequently refer to as the calculated rhabdomeric response.

In the case of axonal responses, we measured spectrally opponent waveforms in all photoreceptor types (Figure 2F-I). Each photoreceptor type exhibits a narrowed tuning in its activation range compared to its calculated rhabdomeric response. In addition, we measured inhibition in the wavelength range outside of the opsin sensitivity. pR7s outputs are activated by UV spanning 320-420 nm, and inhibited by longer wavelengths (Figure 2F, J). yR7 outputs are also activated by UV, with their response remaining excitatory up to 440 nm, and becoming inhibitory from 480 nm onwards (Figure 2G, K). pR8 outputs are the only ones to show a tri-lobed spectral tuning (Figure 2H, L). They are activated by blue light ranging from 400-500 nm and inhibited in the UV from 320-380 nm, as well as in the green from 530-620 nm. yR8s outputs are activated by wavelengths covering the wide range of 400-620 nm in the blue/green but inhibited by UV from 320-380 nm (Figure 2I, M).

Each R7 and R8 terminal type thus displays distinct and specific wavelength opponent properties that, as expected, are dependent on interactions between photoreceptors with different spectral sensitivities. This is generally consistent with previous work. Schnaitmann et al. (2018) found that opponent signals at the level of R7 and R8 outputs are generated through both direct and indirect antagonistic interactions between pairs of R7 and R8 photoreceptors from a single ommatidium: direct interactions in the form of reciprocal histaminergic inhibition, and indirect, inhibitory interaction mediated by a yet-to-be-identified medulla interneuron. However, we measured opponency in ranges that are not predicted by reciprocal inhibition exclusively between R7 and R8 photoreceptors from the same ommatidium. This is most obvious in the case of pR7 and pR8. Indeed, both of these photoreceptor types are inhibited by green light (*>*540 nm) (Figure 2F, H), whereas our measurements of their putative rhabdomeric responses show that neither responds at these long wavelengths (Figure 2B, D). Intra-ommatidial interactions (between R7 and R8 from the same ommatidium) alone are therefore not sufficient to explain the spectral tuning properties we measure. We next aimed to further define the circuit mechanisms that combine and process R7 and R8 rhabdomeric signals to give rise to the diverse spectrally opponent axonal responses that we measured. Our goal was to characterize the effect of these interactions on the spectral encoding properties of photoreceptor outputs and to provide a mathematical description of these interactions.

### Both intra- and inter-ommatidial antagonistic interactions shape the spectral tuning properties of R7 and R8 outputs

According to our rhabdomeric measurements (Figure 2B-E), the inhibition measured in pR7 and pR8s axons in the long wavelength range can only originate from yR8, or the broadband photoreceptors R1-6. Thus, we hypothesized that inter-ommatidial interactions (between R7s and R8s from neighboring ommatidia) and/or inputs from R1-6 contribute to the spectral tuning of R7 and R8 outputs. We employed genetic methods to determine the contribution of specific photoreceptor types to the spectral tuning of R7 and R8 outputs. We took advantage of *norpa-* mutants and selectively rescued NorpA in pairwise combinations of photoreceptor types. We made all possible dual combinations of photoreceptor type rescues, while imaging p/y R7/R8 outputs (Figure 3 and Figure S2).

**Figure 3.**
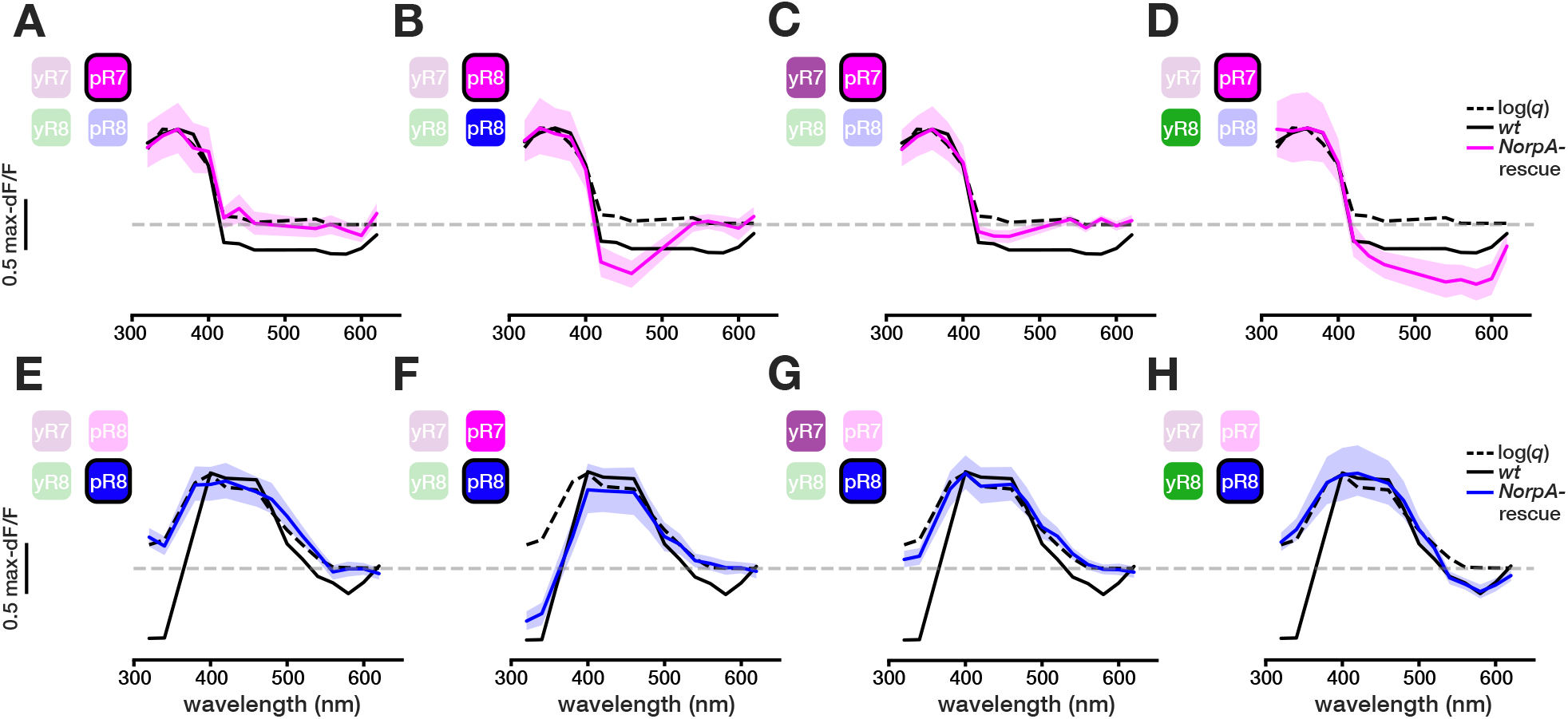
Pairwise NorpA rescues highlight sources of opponency in R7/R8. NorpA, a component of the phototransduction cascade, was restored in *norpa-* blind flies in select pairs of photoreceptor types to determine contributions to opponency. **A-D.** Max-normalized responses of pR7 axons were measured across simulated wavelengths, with NorpA restored in pR7 alone, or in pR7 and a second indicated photoreceptor type. Ns= 106 ROIs (8 flies), 108(8), 132(8), and 104(6), respectively. Dashed black lines represent log(*q*), black lines represent the wild type response, colored lines represent the mean photoreceptor response, shaded regions represents the 95% confidence interval, dashed grey lines represent baseline fluorescence. **E-H.** Maxnormalized responses of pR8 axons were measured across simulated wavelengths, with NorpA restored in pR8 alone, or in pR8 and a second indicated photoreceptor type. Ns= 63(7), 80(9), 69(7), and 63(7), respectively.

First, we imaged pR7 in flies in which pR7 function was restored in combination with one other photoreceptor subtype. The tuning curve of pR7 in flies when phototransduction is rescued in both pR7 and pR8 is similar to that of wild type pR7 in that there is activation in the UV range (320-400 nm) and inhibition in the blue range (420-460 nm) (Figure 3B). This is consistent with intra-ommatidial inhibition from pR8, as the rhabdomeric responses of pR8 show blue sensitivity (Figure 2D). However, inhibition is lost in the long wavelengths (*>*540 nm). In contrast, in a pR7/yR8 rescue, the tuning curve for pR7 displays clear inhibitory responses at all wavelengths above 420 nm (Figure 3D), showing that yR8 contributes to blue/green inhibition in pR7 through inter-ommatidial interactions. In a pR7/yR7 rescue, pR7s are inhibited in the UV/blue range (400-450 nm) (Figure 3C), showing that yR7s contribute to pR7 responses. These results demonstrate that, in addition to intra-ommatidial interactions from pR8, inter-ommatidial interaction from both yR8 and yR7s contribute to opponent responses measured in pR7.

We performed the same set of experiments while imaging pR8 terminals. In a pR8/pR7 rescue, the tuning curve of pR8 becomes bi-lobed, showing inhibition only in the UV range (*<*360 nm) and not in the green wavelength range (*>*540 nm) (Figure 3F). Conversely, in a pR8/yR8 rescue, pR8 still shows inhibition to green but not to UV (Figure 3H). In a pR8/yR7 rescue, we did not see strictly inhibitory responses under our recording conditions, but we did observe a statistically significant decreased response in pR8 in the UV range (300-340 nm) in comparison to the calculated rhabomeric response (Figure 3G). This indicates that yR7 has an inhibitory effect on pR8. These results show that, similarly to pR7s, both intra-ommatidial and inter-ommatidial interactions contribute to the opponent responses measured in pR8s.

We next measured responses in all rescue combinations for yR7 and yR8. In experiments where yR7 was imaged, we confirmed antagonistic inputs from its intra-ommatidial partner yR8 in the green range (*>*540 nm) but could not detect significant inhibition from pR8 or pR7 (Figure S2A-D). yR8 imaging confirmed antagonistic inputs from both p and y R7s in the UV range (*<*380 and *<*400, respectively), but no significant inhibition from pR8 was detected (Figure S2E-H).

Lastly, we investigated the possible contribution of R1-6 to the wild type signals by rescuing NorpA in each photoreceptor type separately together with R1-6. We found no significant differences in paired rescues with R1-6 compared to the measured putative rhabdomeric responses (Figure S2I-P), and thus did not consider R1-6 contributions further in our analysis.

Together, these experiments demonstrate that inter-ommatidial interactions contribute to the spectrally opponent signals that we measure at the output of R7 and R8 photoreceptors, in addition to the intra-ommatidial interactions previously identified. Therefore, inhibitory interactions between R7 and R8 are not confined within medulla columns. Rather, there is a larger set of interactions between columns in the medulla that shape the tuning of R7 and R8 outputs, adding both additional spectral comparisons and a spatial dimension to opponent pathways.

### The horizontal cell-like Dm9 neuron mediates lateral, indirect opponency

The fact that opponent responses in R7s and R8s are shaped by inhibitory interactions between pale and yellow ommatidia is reminiscent of the circuit architecture of vertebrates, where horizontal cells mediate center-surround inhibitory interactions (Thoreson and Mangel, 2012). We thus hypothesized that inter-ommatidial inhibition in the fly medulla is similarly mediated by a horizontal cell-like interneuron in the circuit. The medulla interneuron in question should fulfill the following requirements: 1. be both pre- and post-synaptic to p/yR7s and p/yR8s, 2. span multiple columns (*≥*2) in the medulla, and 3. be excitatory to enable opponent interactions. Only one such neuron has been put forward by electron microscopy (EM) and RNAseq studies: Dm9 (Figure 4A) (Davis et al., 2018; Reiser et al., 2019; Takemura et al., 2013).

**Figure 4.**
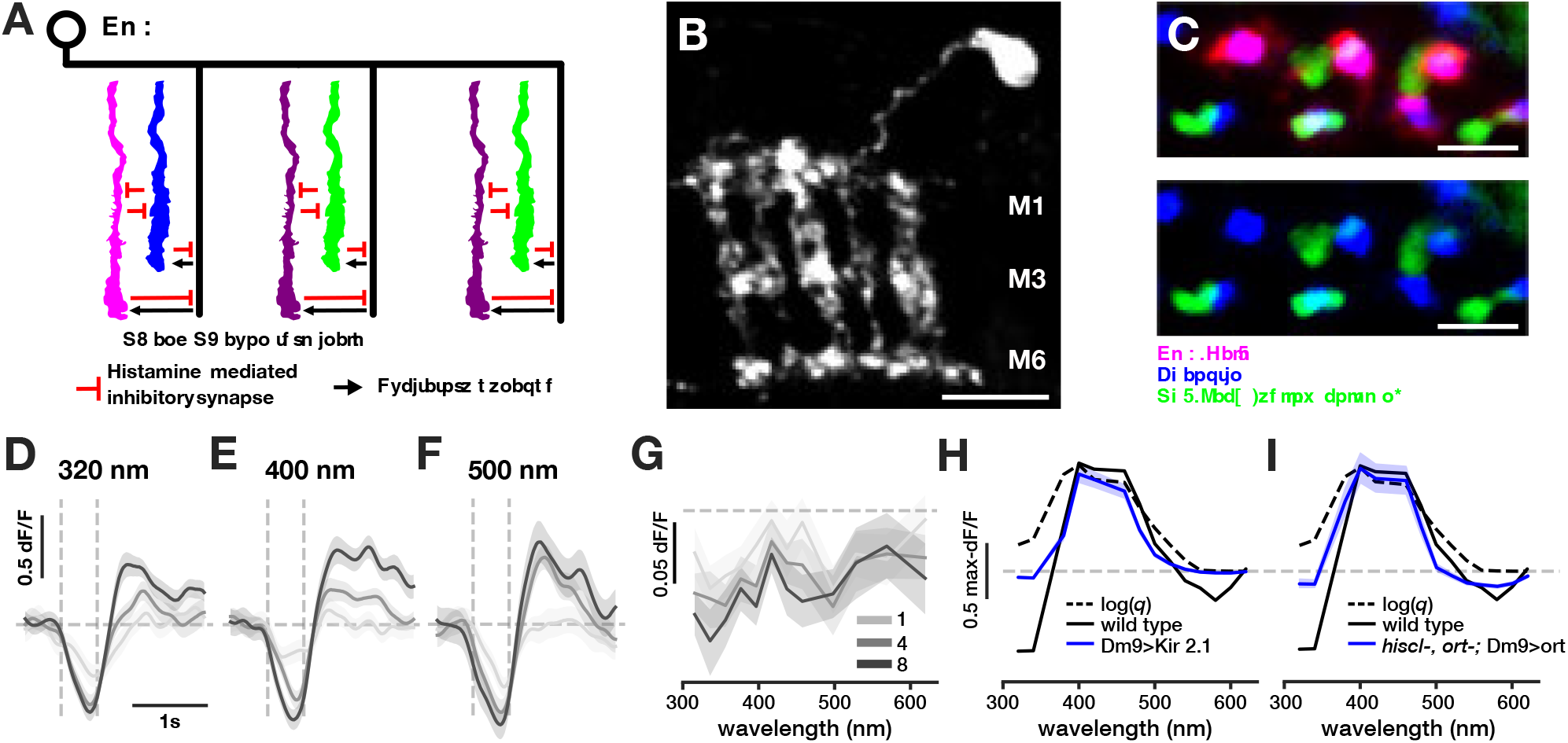
The horizontal cell-like interneuron Dm9 mediates indirect spectral opponency. **A.** Schematic of Dm9/photoreceptor connectivity. Dm9 is an excitatory interneuron spanning multiple medulla columns shown to be both pre- and postsynaptic to R7/R8. **B.** Side view of a maximum projection of a single Dm9 clone (R32E04-Gal4). Scale bar: 10*µm* **C.** Cross section view of a single Dm9 clone (pink), photoreceptor terminals (blue), and yR7 terminals (green) shows a single Dm9 contacts both yellow and pale ommatidia. Scale bar: 5*µm* **D-F.** Responses of Dm9 (R32E04-Gal4) to 0.5 second flashes of three simulated wavelengths over a 10*µE* background with a flat spectrum. Responses to three luminant multiples of each wavelength are shown (1x, 4x, and 8x). Solid lines represent the mean, shaded region represents 95% confidence interval. Vertical dashed grey lines represent onset and offset of light presentation. Horizontal dashed grey lines represent baseline fluorescence **G.** Dm9 spectral tuning curves corresponding to three luminant multiples of each wavelength are shown (1x, 4x, and 8x). **H.** pR8 max-normalized spectral tuning curves. Colored line represents pR8 responses in a Dm9-silenced background (R32E04-Gal4 driving UAS-Kir2.1) N= 323 ROIs (6 flies). Dashed black lines represent log(*q*), black lines represent the wild type responses, and the shaded regions represent the 95% confidence interval. **I.** pR8 max-normalized spectral tuning curves. Colored line represents pR8 in a *hiscl-*,*ort-* mutant background where Ort was rescued in Dm9 (R21A12-GaL4 driving UAS-Ort) N= 153 (6).

Dm9 is a multi-columnar medulla neuron, spanning an average of seven columns and occupying distal medulla layers M1-M6 (Figure 4B). These cells tile in layers M2-M5 but overlap in M1 and M6 (Nern et al., 2015). The processes of Dm9 are in close proximity with R7 and R8 axons in these distal layers of the medulla (Figure S3A-B). EM reconstructions show a large number of synapses from R7 to Dm9 and from R8 to Dm9. EM reconstructions also show synapses from Dm9 back to both R7 and R8. Aside from R7/R8 themselves, Dm9 is the main output of these photoreceptors and also constitutes 50% of the inputs back onto both photoreceptor types. In addition to R7 and R8, Dm9s receive indirect inputs from R1-6 through the lamina monopolar cell L3, as well as inputs from the amacrine cell Dm8 (Takemura et al., 2013). Dm9 has been proposed to be a glutamatergic neuron (Davis et al., 2018). Glutamate in flies can be both excitatory or inhibitory depending on the post-synaptically expressed receptor. The only known fly inhibitory glutamate receptor is GluCl1, which is not expressed in R7 or R8 (Davis et al., 2018). These instead express at least one ionotropic glutamate receptor Ekar, as well as CG11155, predicted to encode another ionotropic glutamate receptor.

Given these properties, we hypothesized that Dm9 is the neuron responsible for indirect antagonistic interactions at the level of R7 and R8 outputs (both intra- and interommatidial). To test this hypothesis, we first measured the spectral tuning of Dm9. We found that Dm9 is inhibited by a broad range of wavelengths spanning the whole spectrum (Figure 4D-G). This is consistent with EM data showing that Dm9 gets inputs from all photoreceptor types. In addition to inhibition to light ON, Dm9 responds positively at light OFF, especially at high intensities of the stimulus. The origin of this OFF response is unclear, but could be due to L3 (Fisher et al., 2015).

Next, we silenced the activity of Dm9 by expressing the inward-rectifying potassium channel Kir2.1 in these neurons specifically (using two different Gal4 lines driving expression in Dm9; Figure S3A-B), while imaging from pR8 axons. We chose this particular photoreceptor type because it provides the clearest read-out of the effect of intra- or inter-ommatidial interactions. UV inhibition in pR8 is likely a combination of intra- and inter-ommatidial interactions, while long wavelength inhibition is due to inter-ommatidial interactions only. We therefore expected only a partial loss in UV opponency after Dm9 silencing, since direct intra-ommatidial inputs from pR7 should not be affected. Conversely, we expected complete loss of inhibition at the long wavelengths with complete Dm9 silencing, as we have shown that the source of these signals is purely inter-ommatidal.

We found that when Dm9 activity is inhibited, the tuning of pR8 is indeed affected (Figure 4H; S3C). Inhibition is overall reduced compared to the spectral tuning in wild type flies, and the tuning of pR8 is no longer tri-lobed. These terminals show opponency in the UV range (300-340 nm) compared to the calculated rhabdomeric response. However, opponency is lost in the green wavelength range (*>*500 nm). This result is consistent with Dm9 mediating inter-ommatidial interactions.

In addition to these silencing experiments, we tested the role of Dm9 in this circuit by disrupting feed-forward inhibition from photoreceptor to Dm9 specifically. Schnaitmann et al. (2018) showed that direct axo-axonal inhibition is mediated by the histamine receptor HisCl1. RNAseq data confirms that HisCl1 is only expressed in photoreceptors, whereas the histamine receptor Ort is expressed in medulla neurons and not in photoreceptors (Davis et al., 2018). Therefore, histaminergic transmission to medulla neurons (including Dm9) is mediated by Ort. As expected, in a *ort-*, *hiscl1-* double mutant background we could not detect any inhibition in pR8 photoreceptors (Figure S3D). We then rescued ort expression exclusively in Dm9 neurons in this mutant background (Figure 4I). When imaging pR8 in these conditions, we found restored opponent waveforms both in the UV range (300-340 nm) and the green range (*>*500 nm), showing that Dm9 is sufficient for mediating interommatidial antagonism.

Our data combined with known connectivity indicates that the horizontal cell Dm9 mediates indirect intra- and inter-ommatidial inhibitory interactions.

### Opponent mechanisms at the level of R7 and R8 optimally decorrelate opsin sensitivities

Our data shows that inter-ommatidial antagonism, in addition to previously described intra-ommatidial antagonism, shapes the responses of the outputs of R7 and R8 photoreceptors. Additionally, we defined the circuit underlying the previously unidentified lateral interactions. What are the consequences of this dual circuit on spectral encoding? To answer this question, we performed comparisons of both calculated rhabdomeric responses and measured axonal responses. In order to do this analysis, we obtained a new dataset of spectral tuning measurements, in which measurements for all photoreceptor types were made with the same stimulus over a large range of intensities. We used a background with a flat spectrum at an intensity of 10*µE* with luminant multiples ranging over several orders of magnitude (Figure S1). The opponent waveforms we measure under these conditions are consistent with our previous experiments (Figure S4).

A clear consequence of opponency is a narrowing of the tuning of the responses in the medulla compared with their rhabdomeric responses (Figure 5A, B). To quantify this, we calculated correlation coefficients between the calculated rhabdomeric responses of R7 and R8 photoreceptors (Figure 5C) and between their measured axonal responses (Figure 5D). As expected from the high degree of overlap between the spectral sensitivities of the four opsins expressed in R7 and R8 (Figure 1D), there is a high degree of correlation between the calculated rhabdomeric responses. This effect is particularly pronounced for spectrally consecutive opsins such as Rh3 and Rh4 (0.97), Rh4 and Rh5 (0.82), and Rh5 and Rh6 (0.79). However, after antagonistic interactions have occurred in the medulla, we find that axonal responses of the different photoreceptor types become decorrelated (yR8 and pR8, yR7 and pR8) and in some cases, anti-correlated (pR7 and both R8s, yR7 and yR8).

**Figure 5.**
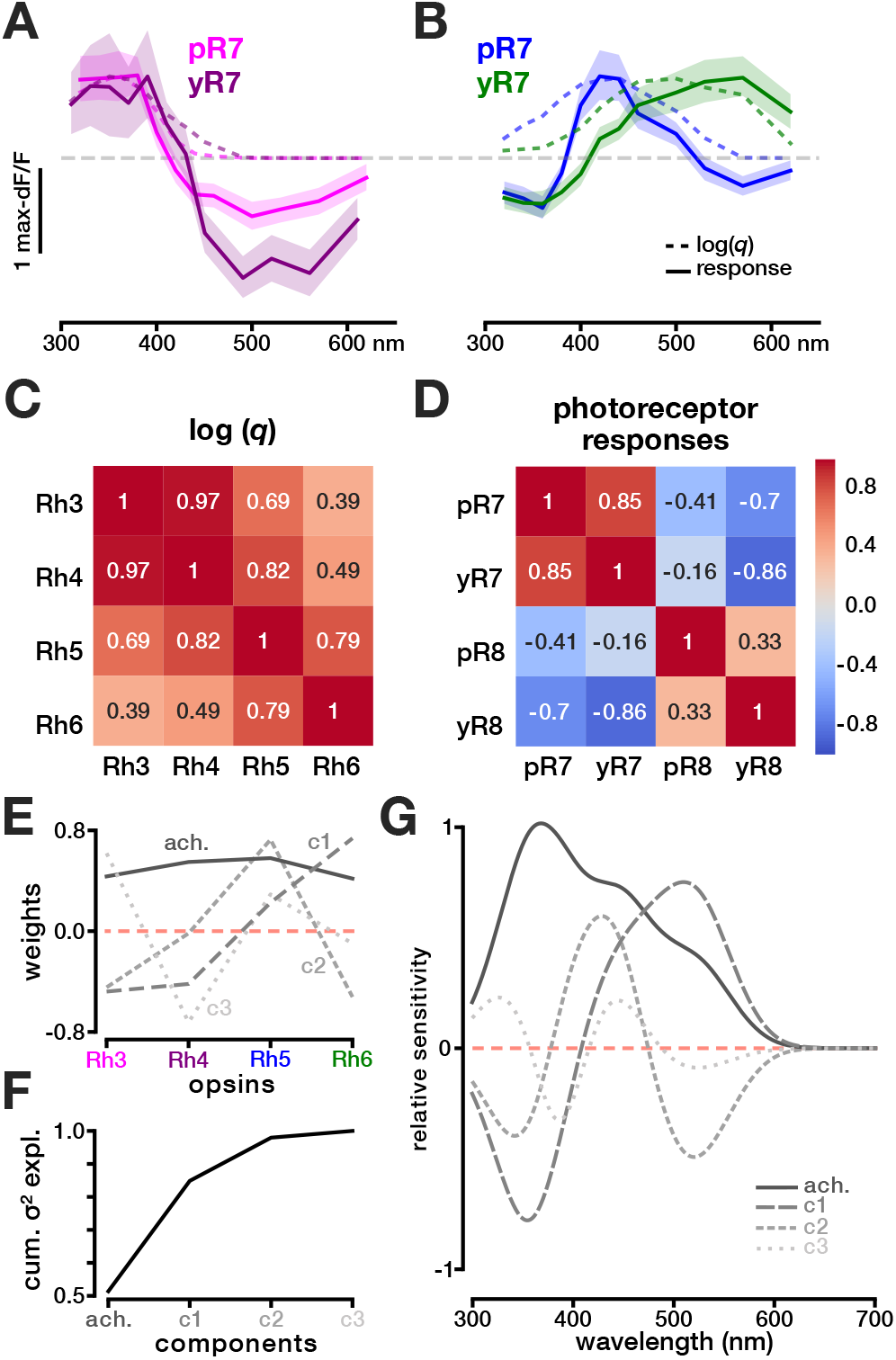
Opponency optimally decorrelates signals along principal components. **A-B.** Comparisons between max-normalized tuning curves based on the calculated rhabdomeric responses (dashed lines) and the measured tuning curves in R7 and R8 axons (solid lines) for the luminant multiple 4 in the flat background (Figure S4). **C.** Correlation matrix comparing the calculated rhabdomeric responses of R7s and R8s. **D.** Correlation matrix comparing the measured axonal responses in R7 and R8 outputs. **E.** Decomposition of opsin spectral sensitivities using principal component analysis (PCA) yields four main principal components. **F.** Percentage of the variance explained by each principal component. **G.** Spectral sensitivity of PCA transformed channels.

The responses we measured at the level of photoreceptor outputs vary along two main axes: one that compares UV and visible wavelengths (y/pR7s and yR8), and one that compares blue with UV + green wavelengths (Figure 5A, B). We asked whether these two axes of opponency were optimal for decorrelating the *Drosophila* opsin sensitivities, or whether other types of comparison would be better suited. Inspired by the Buchsbaum and Gottschalk (1983) study in humans, we decomposed the spectral sensitivities of *Drosophila* R7 and R8 opsins using principal component analysis (PCA) (Figure 5E-G). The first principal component (PC) is achromatic, with equal loading for all opsin types, and it accounts for over half of the variance. Higher PCs therefore describe variance in the chromatic domain. The second PC opposes the two R7 opsins and the two R8 opsins, corresponding to comparison between the UV and the visible parts of the spectrum. The third PC opposes Rh5 and Rh3 + Rh6, which corresponds to a comparison between blue and UV + green. The last PC opposes Rh3+Rh5 and Rh4+Rh6. The first two chromatic PCs together with the achromatic PC explain 97% of the variance. Interestingly, these two chromatic PCs broadly describe the two types of responses we measure at the output of R7 and R8: UV vs visible (observed in pR7/yR7 and yR8) and blue vs UV + green (observed in pR8). The first chromatic axis is supported by intra-ommatidial interactions, whereas the second chromatic axis necessitates inter-ommatidial interactions. The opponent responses encoded by the terminals of these photoreceptors are thus consistent with an optimal decomposition of the spectral sensitivities of the opsins they express.

### A recurrent model of early color circuits predicts spectrally opponent R7 and R8 outputs

We next asked whether the circuit architecture we identified and tested experimentally can quantitatively reproduce the opponent responses we measure in R7 and R8 outputs. To inform the construction of our model, we first tested the linearity of the system. Specifically, we asked if photoreceptor axonal outputs are linear with regard to their rhabdomeric inputs (i.e. log(*q*)). To test for linearity, we assessed two empirical measures: scalar invariance and additivity. The estimated zero-crossing points of the opponent tuning curves of R7s/R8s do not significantly change at different intensities of light measurements, showing scalar invariance within the bounds of our recording conditions (Figure S5A). To test for additivity, we measured the responses of photoreceptor outputs to wavelengths mixed in different ratios (see methods; Equation 8) and compared the responses to corresponding linear additions of the single wavelength responses. Figure S5B-F shows the results for mixtures between 340/440 nm, 380/620 nm, 400/570 nm, 460/570 nm, and 320/530 nm at four different mixing ratios. The measured responses to the mixtures (filled circles) do not significantly differ from the linear predictions (shaded area). Therefore, the circuit under investigation behaves linearly within the range of stimuli used in this study.

Because photoreceptors integrate rhabdomeric inputs linearly, we first performed a linear regression without biological constraints. We used the calculated rhabdomeric responses as independent variables to fit our amplitude measurements in the flat background condition (see methods; Equation 13). We found that comparisons between measured axonal responses and estimated responses based on linear regression fall on the unity line (Figure S5G-J), thus providing a good fit (Figure 6A). The unconstrained linear regression provides a benchmark for our next model which includes biological circuit constraints.

**Figure 6.**
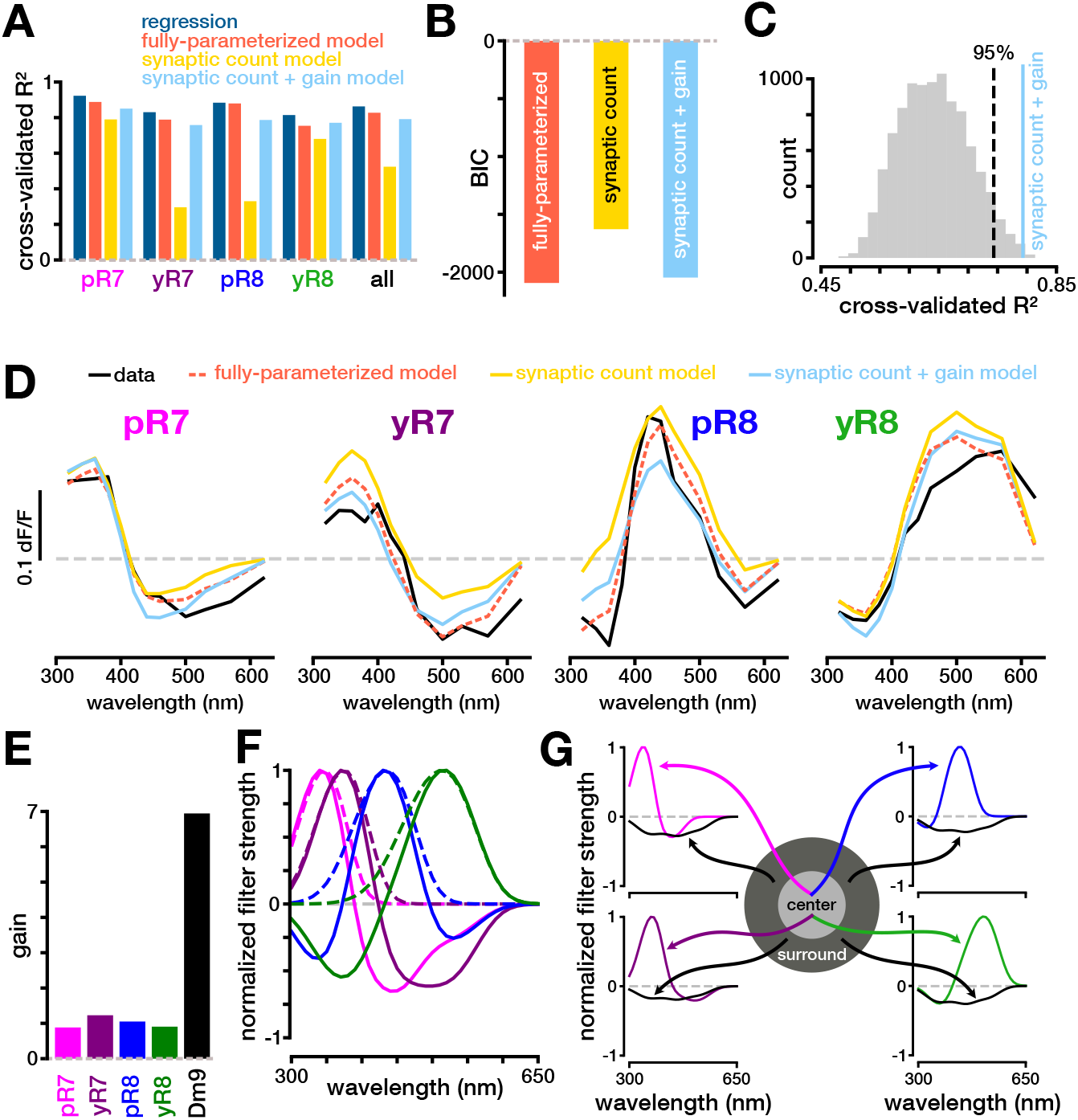
Recurrent model of color opponency in R7 and R8 photoreceptors. **A.** Comparison of different cross-validated *R*2 values using linear regression, the fully-parameterized recurrent model, the synaptic count recurrent model, and the synaptic count + gain recurrent model. **B.** Comparison of BIC values for the different iterations of the recurrent model. **C.** Distribution of *R*^2^ values using random weights for the synaptic count + gain model. The dotted line indicates the 95th percentile of the distribution and the solid colored line indicates the *R*^2^ value using the synaptic counts as weights. **D.** Predicted responses for the different iterations of the recurrent model (colored), and the actual mean response of the photoreceptor in question (black). **E.** Fitted gains for different neurons in the recurrent circuit for the synaptic count + gain model. **F.** The predicted spectral filtering properties of the different photoreceptor outputs (solid line) compared to the filtering properties of the rhodopsin they express (dashed line). **G.** The spectral filtering properties for the predicted center and surround of the different photoreceptor outputs.

We next built a linear recurrent network constrained by the connectivity and signs within this network (see methods; Equation 14). The overall architecture of the network consists of direct inhibitory connections between photoreceptors within a single ommatidium, and indirect connections via the excitatory interneuron Dm9. Dm9 receives inhibitory inputs from all four photoreceptors and feeds back onto all photoreceptors. We fit the steady state of this model to our measured amplitudes (see methods; Equation 16). When constrained by this architecture, our model has goodness-of-fits comparable to the unconstrained linear regression, even though the fully-parameterized recurrent model uses fewer parameters (Figure 6A). Our model also provides fitted tuning curves that closely approximate the opponency we observe in our data (Figure 6D). This shows that our proposed circuit is mathematically plausible given our measured responses, while also providing a biological constraint consistent with our experimental observations.

Finally, we aimed at further constraining the model by using synaptic counts obtained by EM reconstructions as a proxy for synaptic weights (Reiser et al., 2019; Takemura et al., 2015). To compare the performance of the different models, we calculated the Bayesian information criterion (BIC) (see methods; Equation 9), which provides a metric that balances the overall error with the number of parameters in the model (a lower BIC value is preferred). The “parameter-free” model, which we refer to as the synaptic count model, qualitatively predicts our data, as seen in the tuning curves it produces (Figure 6D). However, it does not quantitatively perform as well as the fully parameterized model (Figure 6A, B). This is likely due to the fact that in this synaptic count model, we make the explicit assumption that the gains of the different neurons in the circuit are equal, which is not necessarily biologically plausible. We therefore fitted our data to a model that includes fitted gain parameters for each of the photoreceptors and Dm9 separately, keeping synaptic counts as synaptic weights for the connections between neurons. We refer to this as the synaptic count + gain model. The gain parameters we obtain are similar between photoreceptor types, and larger for Dm9 (Figure 6E). This model performs just as well as the fully parameterized model, both qualitatively and quantitatively (Figure 6A, B, D). As a control, we replaced the weights in our model with randomly drawn sample weights 10,000 times, and created a distribution of *R*^2^ values (Figure 6C). We found that using the synaptic count for our weights results in a significantly better performance than when using random weights. Therefore, the synaptic count data retrieved from EM gives a non-random estimate of input strength to photoreceptors.

We used the synaptic count + gain model to predict the spectral sensitivities of photoreceptor outputs, not only to a full-field stimulus, corresponding to our experimental paradigm, but also to spectrally varying stimuli in the center and the Dm9-mediated surround separately. In Figure 6F, we used our model to predict the full-field spectral filtering properties of each photoreceptor. These filtering properties reflect our experimental tuning curves, but also predict the response of photoreceptors to arbitrary spectral distributions. In Figure 6G, we modeled the sensitivities of the center and the surround separately. We found that sensitivities in the center are in all cases bi-lobed, corresponding to comparisons between the UV part of the spectrum and the visible part, as expected. Additionally, we found that the sensitivities of the surround are broadband and strongest between 350 nm and 500 nm. The predictions made by our model lay the groundwork for future experiments in which spatially patterned stimuli can be used to further explore how this circuit processes information both spatially and spectrally.

## Discussion

In this work, we report the spectral tuning of wavelength-specific R7 and R8 photoreceptor outputs in the visual system of the fruit fly *Drosophila melanogaster*. We find that each R7 and R8 output displays distinct spectrally opponent properties to full field presentations of light. p and yR7 are both activated by UV light and inhibited by visible light, with shifted sensitivities. pR8s are activated by blue and inhibited by both UV and long wavelength green light. yR8s are activated by visible wavelengths and inhibited by UV. These signals are a consequence of a dual circuit: one that consists of reciprocal inhibition between R7 and R8 from the same ommatidum, and another that supports lateral inhibitory interactions between R7s and R8s from neighboring ommatidia. We show that the latter is mediated by the horizontal-cell-like Dm9 neuron, which both gets inputs from and feeds back onto all R7 and R8 photoreceptor types. Interestingly, a consequence of this dual circuit is optimal decorrelation of photoreceptor signals. We built an anatomically constrained linear recurrent model which describes our findings and shows that synaptic count is a quantitative predictor of circuit function. We also predict the spatio-chromatic receptive field structure of each photoreceptor using our mathematical model.

### Both intra- and inter-ommatidial antagonistic interactions contribute to the spectral tuning of R7 and R8 photoreceptors

At the circuit level, the spectral tuning properties of R7 and R8 outputs are a consequence of the superposition of two types of antagonistic interactions: reciprocal inhibition between R7 and R8 of the same ommatidial-type (Schnaitmann et al., 2018), enabling UV vs visible comparison at one point in space, and inter-ommatidial interactions between R7 and R8 from neighboring ommatidia, allowing for additional comparisons to be made in the spectral domain (e.g. blue vs green) by comparing photon catches at different points in space.

We found that the indirect interactions are mediated by the interneuron Dm9. EM describes the connectivity of Dm9 as both pre- and postsynaptic to photoreceptors, making it the equivalent of horizontal cells in the mammalian retina. Its main direct inputs are from both p and y R7s and R8s. Based on synaptic count, Dm9 also receives a small number of indirect inputs from R1-6 through the lamina monopolar cell L3, and even fewer inputs from the amacrine-like neuron Dm8, a postsynaptic partner of R7 photoreceptors. According to this connectivity, we were expecting to find functional evidence for inhibition from all R7/R8s as well as from R1-6 onto all R7/R8s. However, under our recording conditions, we could only measure a subset of these. All the combinations where we could not detect inhibition correspond to interactions from p onto y subtypes (pR7 onto yR7, pR8 onto yR8 and pR8 onto yR7). The only such interaction that we could detect was from pR7 onto yR8, which express opsins with the most disparate spectral sensitivity. We hypothesize that the lack of detectable inhibition is likely a combined effect of the substantial overlap of the opsin sensitivities in the range of our measurements, and the lower ratio of p vs y ommatidia in the fly eye, together leading to activation of the imaged photoreceptor overcoming inhibition from surrounding photoreceptors in our measurements. In addition, we could not detect a contribution of R1-6 to the responses of R7/R8s. This could be due to the same issue of overlap, combined with the relatively smaller strength of R1-6 inputs onto Dm9. It has also been shown that there are gap junction between R1-6 and R8 (Wardill et al., 2012), which may mitigate the inhibitory effect of R1-6 photoreceptors onto R8. It is possible that the stimulus composition will affect these functional interactions between photoreceptor types and that in some condition of illumination these interactions contribute more to the outputs of R7 and R8.

### Spectral opponency optimally decorrelates photoreceptor output signals

Theories of efficient coding postulate that the purpose of the early visual system is to compress redundant information and remove noise prior to neural transmission (Atick and Redlich, 1992; Barlow, 1961). Redundancy stems from correlations that occur extrinsically, in the statistics of natural scenes (chromatically, spatially and temporally), but also intrinsically, produced by the strong spectral overlap of photoreceptors’ opsin sensitivities. One well-known way of removing these correlations is via a linear decomposition (Atick and Redlich, 1992; Barlow, 1961; Buchsbaum and Gottschalk, 1983). In accordance with this, we showed that the photoreceptor outputs of *Drosophila* perform a linear transformation on the inputs that orthogonalizes photoreceptor responses (decorrelation) and creates opposing, near-symmetric chromatic channels (strong anti-correlation). pR7/yR7s and yR8s compare the UV vs the visible part of the spectrum (all crossing over between 380-430 nm), forming near-mirror images of each other. pR8s are the only photoreceptors with a three-lobed sensitivity, comparing blue to both UV and green. Using PCA analysis, we found that these axes of opponency, which are a result of both intra- and inter-ommatidial antagonism, are optimally suited to decorrelate opsins sensitivities - a feature shared with trichromatic primate retinas (Buchsbaum and Gottschalk, 1983). However, spectral decorrelation is likely not the only goal, as pR7, yR7, and yR8 photoreceptors all encode spectral inputs along the UV-vs-visible axes. The absolute value of the correlation coefficients of pR7-yR8 and yR7-yR8 outputs are actually larger than the absolute value of their predicted correlation coefficients at the level of the retina. Such redundancy may serve to deal with noise in the system, so that visual stimuli along the UV-vs-visible axis can be robustly encoded. This circuitry could effectively support behaviors that depend on differences between short and long wavelengths.

### A circuit constrained recurrent model predicts R7 and R8 spectrally opponent outputs with complex spatio-chromatic receptive fields

By building an anatomically constrained model of the underlying circuit, we showed that the circuit architecture we identify can quantitatively produce the signals we measure at the level of R7 and R8 outputs. We further constrained our model using synaptic counts obtained from EM. Synaptic count data has been previously used to gain intuition about which inputs to a given neuron are likely strongest (Scheffer and Meinertzhagen, 2019). However, it was not clear whether synaptic count could be used more quantitatively to predict function. Using our model, we showed that synaptic count is both a good qualitative and quantitative estimate of synaptic strength. This result demonstrates that, at least in this type of hardwired sensory circuit, synaptic count provides useful information for understanding circuit function.

We used our biologically constrained model to make predictions of the responses of photoreceptor outputs to untested visual stimuli. Our model predicts that the result of this dual opponent system is a spatio-chromatic receptive field for each photoreceptor output with a UV vs visible color-opponent center and an antagonistic achromatic surround (Figure 6F). The size of the center is predicted to correspond to one ommatidial angle, or *∼*5 degrees. The size of the surround is likely determined by the columnar extent of the horizontal cell Dm9, which has been found to span on average 7 columns (Nern et al., 2015), corresponding to 35 total degrees in visual space (with a width of *∼*15 degrees). As in the case of mammalian horizontal cells, electrical synapses may increase receptive field size and possibly be modulated by illumination conditions (Zhang et al., 2013). Due to the limitations of our current visual stimulus system, these receptive fields were not tested experimentally. Moving forward, a patterned chromatic visual stimulus will enable direct measurements of receptive field sizes and spectral properties, and enable testing of the predictions of our model.

### Functional implications

The Dm9-mediated, inter-ommatidial circuit that we describe here can be directly compared to mechanisms that establish opponency in the retina of trichromatic primates. There, midget cells compare photon catches between M and L cones, creating a red-green opponent axis. This opponent channel is thought to be established through non-selective wiring of H1 horizontal cells with M and L cones (Crook et al. 2011; Lennie et al. 1991; but see Lee et al. 2012). In the fovea, each midget ganglion cell receives inputs from a single M or L cone at its center, and a mixture of M and L cones in its surround. This so-called “private line” circuitry supports both high acuity and cone opponency, resulting in multiplexed signals capturing both high-resolution achromatic stimuli that isolate the center and low-spatial resolution chromatic stimuli that engage both center and surround (Atick et al., 1992; Derrico and Buchsbaum, 1991). The ambiguity between these multiplexed signals may be resolved by differential processing into two downstream pathways, which preserve either chromatic signatures at the expense of spatial information, or vice versa. This diversity of processing leads to blobs and inter-blobs in the visual cortex, where chromatic and achromatic information about the visual scene are encoded, respectively (Livingstone and Hubel, 1984; Tootell et al., 1988).

The circuit we describe at the level of photoreceptor outputs is similar to the foveal midget pathway: it is horizontal-cell-mediated, samples the center at one point in space, samples the surround randomly from the distribution of opsins in the eye, and creates spatially and spectrally opponent responses. However, there are two main differences in the fly. First, because the surround samples from both R7s and R8s, its sensitivity is predicted to encompass the whole light spectrum available to the fly independent of its p/y composition. Various p/y compositions would only result in deviations at the shorter and longer tails of the wavelength spectrum. Second, and most importantly, the center itself has spectrally opponent properties, supported by direct axo-axonal synapses between R7 and R8 from the same ommatidium. This type of center-surrround arrangement results in the complex receptive fields we model with a UV vs visible center and an antagonistic broadband surround. R7 and R8 photoreceptor signals thus also convey multiplexed information, which could be differentially processed downstream in a manner analogous to the midget pathway. However, because of the additional spectral opponency in the center of the receptive field, signals in the fruit fly would be separated into a high-resolution chromatic pathway and a low-resolution chromatic pathway. In addition, because they are active in daylight, R1-6 provide inputs to an achromatic high-resolution pathway.

Unlike the simple eye of mammals, the compound eye of the fly is not subject to the limitations of optical aberration. It thus has the capacity to build a chromatic comparison system that operates at the full resolution of the eye, equivalent to the resolution of achromatic pathways. It uses an insect specific circuit architecture that is well suited to extract chromatic information for small target visual stimuli, at a scale equivalent to the resolution of the fly eye (*∼*5 degrees). Additionally, the fly uses a horizontal-cell-mediated circuit based on lateral interactions, similar to the one used in primates. This system allows for further chromatic comparisons to be made, like the one we measured between the blue and green parts of the spectrum. By pooling signals in space, the horizontal cell pathway may increase sensitivity of responses to small chromatic differences. This circuit architecture is well tuned to extract chromatic information for large target visual stimuli. Overall, the dual circuit that exists at the level of R7 and R8 outputs combines an insect specific circuit motif, which could enable chromatic vision at the full resolution of the fly eye, and an evolutionarily convergent center-surround circuit motif, which could allow for lower spatial resolution chromatic vision with extended spectral resolution.

## Acknowledgments

We benefited from helpful conversations with L. Abbott and T. Movshon. We thank L. Abbott, C. Desplan and C. Zuker for comments on the manuscript. We thank M. Reiser for sharing data prior to publication, L. Zipusky for advice on Dm9 Gal4 lines, F. Rouyer, D. Hattori and M. Wernet for fly stocks, D. Peterka for advice on imaging, R. Hormigo for help with the fly back system. RB was supported by NIH R01EY029311, the McKnight Foundation, Grossman Charitable Trust and the Kavli Foundation. SLH acknowledges support from NSF GRF DGE-1644869. JRK was supported by NIH F31EY029592. MPC, EO and MSW and this project were supported by NIH R01EY029311.

## Contributions

RB, SLH, and MPC conceived of experiments and wrote the manuscript. SLH acquired the imaging data. SLH and MSW performed animal husbandry. MPC processed and analyzed the imaging data, wrote the visual stimulaton code, and performed the modeling work. EO built the flyback system. MSW and EO acquired immunohistochemistry data. MSW and JRK genotyped flies.

## Methods

### Genetics

w+ flies were reared on standard molasses-based medium at 25°C - 28°C. The rhodopsin drivers used for imaging photoreceptors Rh3-Gal4 and Rh6-Gal (Cook et al., 2003) along with Rh4-Gal4 and Rh5-Gal4 (Saint-Charles et al., 2016) were expressed heterozygously along with 20X-UAS-GCamp6f, also expressed heterozygously (Bloomington stock center: 52869). Dm9 cells were targeted for imaging, staining and silencing using both the R21A12-Gal4 or the R32E04-Gal4 drivers (Bloomington stock center: 48925 and 49717). Silencing was performed using UAS-Kir2.1 constructs (made and gifted by by Daisuke Hattori), and imaging with Rh5-LexA (Vasiliauskas et al., 2011) (gift from Claude Desplan). Photo-transduction rescue experiments were performed using*norpa-* and UAS-NorpA1 or UAS-NorpA2 constructs (Wernet et al., 2012) (gifts from Mathias Wernet). Ort rescue experiments were performed in a *hiscl*^134^*ort*^1^ background citep (gift from Mathias Wernet), heterozygous with *hiscl*^134^*ort*^1^*cry*^02^ (Alejevski et al., 2019) by also expressing a UAS-ort construct (Alejevski et al., 2019) (both gifts from Francois Rouyer). For immunostaining, UAS-mCD8 (gift from Claude Desplan) or UAS-GCaMP6f (Bloomington stock center: BL42747) constructs were used to label cell types of interest. For clones, hs-FLPG5.PEST and 10XUAS(FRT)myr::smGdP-V5/FLAG/HA-10XUAS(FRT) constructs were used (Bloomington Stock center 64085) as well as Rh4-LacZ (Nern et al., 2015) (Gift from Claude Desplan).

### Two-Photon Calcium Imaging

Imaging was conducted with a two-photon microscope (Bruker) controlled by PrairieView 5.4 and a mode-locked, dispersion compensated laser (Spectraphysics) tuned to 930 nm. We imaged with a 20x water-immersion objective (Olympus XLUMPLFLN, 1.0 numerical aperture). In front of the photomultiplier tube (Hamamatsu GaAsP), we mounted a bandpass filter (Semrock 514/30 nm BrightLine) to reduce bleed-through from the visual stimulus setup. T-Series were acquired at 15-30Hz and lasted for a maximum of eight minutes with each frame at x-y imaging being 145×90 pixels.

All experimental animals for functional imaging were briefly anaesthetized using carbon dioxide on the day of eclosion, and imaged at ages ranging from 3-13 days. Flies were prepared for two-photon imaging based on methods previously described (Behnia et al., 2014). Flies were anesthetized using ice, and mounted in a custom stainless-steel/3D-printed holder. A window was cut in the cuticle on the caudal side of the head to expose the medulla, where the axons of photoreceptors could be imaged. The eyes of the of the fly remained face down under the holder, and remained dry while viewing the visual stimuli, while the upper part of the preparation was covered with saline. The saline composition was as follows (in mM): 103 *NaCl*, 3 *KCl*, 5 *n−tri*(*hydroxymethyl*) *methyl−*1*Aminoethane−sulphonic acid*, 8 *trehalose*, 10 *glucose*, 26 *NaHCO*_3_, 1 *NaH*_2_*PO*_4_, 1.5 *CaCl*_2_, and 4 *MgCl*_2_, adjusted to 270mOsm. The pH of the saline was equilibrated near 7.3 when bubled with 95% *O*_2_ / 5% *CO*_2_ and perfused continuously over the preparation at 2 *ml/min*. The imaging region of interest was limited to the region of the medulla photoreceptors are directly activated by stimuli. The dorsal third of they eye was covered with black acrylic paint. Calcium responses were stable throughout imaging.

### Immunohistochemistry

Immunostainings were done as described by Morante and Desplan (2011) with some modifications. Adult flies were anesthetized on ice. Brains were dissected in PBS and fixed in 4% formaldehyde for 35 minutes on ice. Brains were incubated at 4°C overnight with the following primary antibodies: sheep anti-GFP (1:500, AbD Serotec), rat anti-DN-cadherin (1:50, DSHB) and mouse anti-chaoptin (1:50, DSHB) diluted in PBST (0.3% TritonX-100 in PBS). Secondary antibodies were incubated for 2 hours at room temperature. Images were acquired using an Nikon A1R Confocal Microscope.

To obtain Dm9 clones, 2-3 day old flies were heat shocked for 3 minutes at 3°C and dissected 2 days later. Dm9 clones were labeled with the FLAG epitope tag using the primary antibody rat anti-DYKDDDDK (1:200, NBP1). Yellow R7 photoreceptors were labeled with an Rh4-LacZ reporter construct and the primary antibody rabbit anti-beta-galactosidase (1:2000, MBP).

### Visual Stimulation

#### Hardware

We produced full-field wavelength-specific stimuli using a customized setup (Figure 1C). The setup consists of six LEDs in the UV and visible wavelength range (ThorLabs M340L4 - dUV/340nm; M365L2 - UV/360nm; M415L4 - violet/415nm; M455L3 - blue/455nm; M565L3 - lime/565nm; M617L3 - orange/615nm). A customized driver drove the five LEDs from dUV to lime. These LEDs turned on during the return period of the x-scanning mirror in the two-photon microscope (fly-back stimulation). We used the TTL signal generated by the two-photon microscope at the beginning of each line-scan of the horizontal scanning mirror (x-mirror) to trigger the LED driver. An individual T-Cube (Thorlabs LEDD1B T-Cube) drove the orange LED. Stimuli were generated using customized software written in Python. The update rate for the LED voltage values was 180Hz.

The different light sources were focused with an aspheric condenser lens (ThorLabs ACL2520U-A) and aligned using dichroic mirrors (dUV-UV dichroic - Semrock LPD01-355RU; UV-violet dichroic - Semrock FF414-Di01; violet-blue dichroic - Semrock Di02-R442; blue-lime dichroic - Semrock FF495-Di03; lime-orange dichroic - Semrock FF605-Di02). The collimated light passed through a diffuser (ThorLabs DG10-1500A) before reaching the eye of the fly.

#### Intensity calibration

In order to measure the intensity of our LEDs across many voltage outputs, we used a photo-spectrometer (250-1000 nm, Ocean Optics) that was coupled by an optic fiber and a cosine corrector and was controlled using our customized Python software. The photo-spectrometer was mounted on a 3D printed holder that was designed to fit on our experimental rig and approximately aligned with the fly’s point of view. For each LED, we tested a total of 40 voltage values (linearly separated) from the minimum voltage output to the maximum voltage output. For each voltage value tested, we adjusted the integration time to fit the LED intensity measured, and averaged over 20 reads to remove shot noise.

Using the spectrometer output, we calculated the absolute irradiance (*I*_*p*_(*λ*); in *W/m*^2^*/nm*) across wavelengths using the following equation:

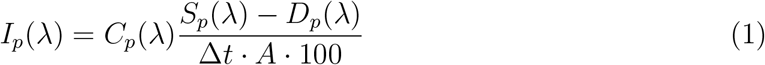

where *C*_*p*_(*λ*) is the calibration data provided by Ocean Optics (*µJ/count*), *S*_*p*_(*λ*) is the sample spectrum (*counts*), *D*_*p*_(*λ*) is the dark spectrum (*counts*), ∆*t* is the integration time (*s*), and *A* is the collection area (*cm*^2^).

**Table S1.**
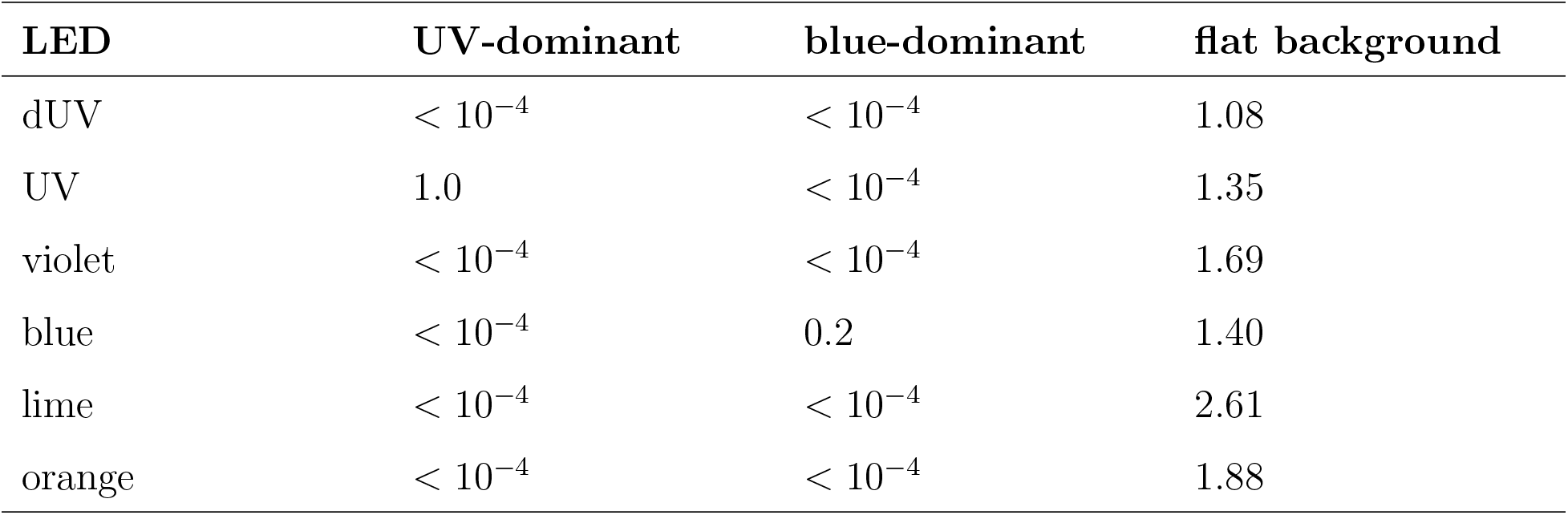
Background light intensities. Different background light intensities used for the different stimulus conditions. Related to Figure 1–6.

| LED | UV-dominant | blue-dominant | flat background |
| --- | --- | --- | --- |
| dUV | $< 10^{-4}$ | $< 10^{-4}$ | 1.08 |
| UV | 1.0 | $< 10^{-4}$ | 1.35 |
| violet | $< 10^{-4}$ | $< 10^{-4}$ | 1.69 |
| blue | $< 10^{-4}$ | 0.2 | 1.40 |
| lime | $< 10^{-4}$ | $< 10^{-4}$ | 2.61 |
| orange | $< 10^{-4}$ | $< 10^{-4}$ | 1.88 |

Next, we converted absolute irradiance to photon flux (*E*_*q*_; in *µE/nm*):

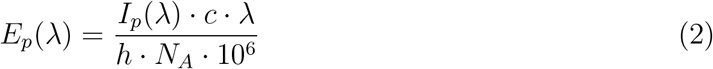

where 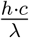 is the energy of a photon with *h* as Planck’s constant (6.63 *⋅* 10^*−*34^ *J ⋅ s*), *c* as the speed of light (2.998 *⋅* 10^8^ *m/s*), and *λ* the wavelength (*nm*). *N*_*A*_ is Avogadro’s number (6.022 *⋅* 10^23^ *mol*^−1^).

#### Stimulus Design

Each stimulation protocol had 10-20 seconds before and after the stimulation period in order to measure baseline fluorescence (fluorescence to background light).

The intensities of each LED for the different background conditions are shown in table S1. Flies were adapted to the differenct background lights for approximately 5 minutes before the start of the recording. For the flat background condition, we chose the intensities of the LEDs by fitting the following equation:

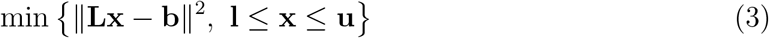

where **L** is a matrix of the normalized LED intensities across wavelengths (each row is a different wavelength and each column is a different LED), **x** is a vector of corresponding LED intensities to fit, and **b** is the background spectrum across wavelengths (i.e. a flat spectrum with an overall intensity of 10 *µE*). **x** is bounded by the minimum **l** and maximum **u** intensity each LED can reach. The minimum intensity is zero for all LEDs, and the maximum intensities are (in *µE*): dUV - 11.7, UV - 21.7, violet - 17.0, blue - 16.4, lime - 18.7, and orange - 145.1.

We wanted to show different single Gaussian wavelengths between 320-620 nm with a standard deviation of 10 nm on top of our background (i.e. add these single wavelengths to our background light) (Figure 1F). We also wanted to show these single wavelengths across different intensities. To do this, we built a simple model of opsin photon capture.

The absolute photon capture of an opsin (i.e. the number of photons absorbed) given any spectral stimulus at a specific intensity can be calculated as follows (Kelber et al., 2003; Renoult et al., 2017):

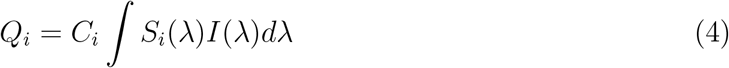

where *Q*_*i*_ is the absolute photon capture of opsin *i*, *C*_*i*_ is the absolute sensitivity of opsin *i*, *S*_*i*_ is the relative spectral sensitivity of opsin *i*, and *I* is the spectrum of light entering the eye. Equation 4 implies that the identity of a photon is lost upon absorption by a photoreceptor (i.e. the principle of univariance). As the scaling factor *C*_*i*_ is usually unknown, the relative photon capture can be calculated instead assuming von Kries chromatic adaptation (Kelber and Osorio, 2010; von Kries, 1904; Renoult et al., 2017):

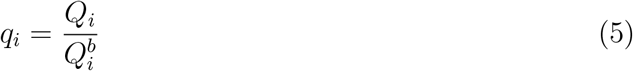

where *q*_*i*_ is the relative photon capture of opsin *i*, and 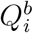 is the absolute photon capture of opsin *i* for the background illuminant.

For our six LEDs, we can calculate the normalized relative capture across the fly opsins:

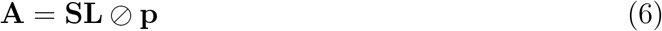

where **A** is a matrix corresponding to the relative photon capture of each opsin for each LED (*opsin × LED*), **S** is a matrix of the relative spectral sensitivities for all opsins across wavelengths (*opsin × wavelength*), **L** is a matrix of the normalized LED intensities across wavelengths (*wavelength × LED*), and **p** is a vector of the absolute capture for all opsins for the background spectrum. ⊘ signifies element-wise division.

To emulate our desired stimuli using our six LEDs, we first calculate the relative photon capture of each opsin present in the fly eye given the desired stimulus. This gives us a vector **q**. Given **A** from equation 6, we find the optimal intensities for each LED to match our desired **q** as follows:

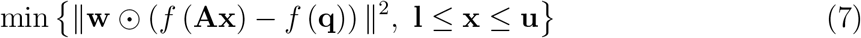

where **x** is a vector of corresponding LED intensities to fit, *w* is a weighting factor for each opsin, and *f* is a link function (i.e. the identity for the single wavelength dominant backgrounds and the log for the flat background). The weighting factor *w* was 1 for all opsins in the case of the single wavelength dominant backgrounds, and 1 for all opsins, except 0.1 for rh1, in the case of the flat background. The lower (**l**) bound on **x** corresponds to the background intensity of each LED, as we desired to add a spectrum on top of the background. The upper (**u**) bound on **x** correspond to the maximum intensity each LED can reach. ⊙ signifies element-wise multiplication.

We used a total of three stimulus sets. The accuracy of our fitting procedure is shown in Figure S1G-R. Each individual stimulus (i.e. each simulated wavelength or wavelength mixture) lasted 0.5 seconds with a 1.5 second period between stimuli. The background intensity values are shown in Table S1. Our UV-dominant and blue-dominant background was used to test the existence of color opponency in R7s and R8s, respectively. Both stimulus sets had a total of 16 wavelengths that were tested spanning 320 to 620 nm, and each stimulus was repeated three times. In the case of the UV-dominant background, each wavelength was fitted using an intensity that was 5 times bigger than the total background intensity (i.e. a luminant multiple of 5). In the case of the blue-dominant background, the wavelengths were a luminant multiple of 15. In the case of the UV-dominant background, we discarded the simulated wavelengths 480, 500, and 520 nm, because the dUV LED is on for these longer fitted simulated wavelengths; the algorithm was trying to fit the relative capture of the broadband rh1 opsin (Figure S1A, G, M). In the case of the blue-dominant background, we discarded the wavelengths 360 and 440 nm, because the green and orange LED is on respectively for these shorter fitted simulated wavelengths; the algorithm was trying to fit the relative capture of the broadband rh1 opsin (Figure 1B, H, N). To avoid this issue during fitting of the flat background stimuli, the error for the rh1 capture is weighted differently (Equation 7). This is reasonable considering R1-6 photoreceptors do not contribute significantly to R7 and R8 photoreceptor responses (Figure S2M-Q).

For the flat background our single wavelengths included: 320 nm, 340 nm, 360 nm, 380 nm, 400 nm, 420 nm, 440 nm, 460 nm, 500 nm, 530 nm, 570 nm, 620 nm. We tested luminant multiples of 0.2, 1, 4, and 8. We also mixed the wavelengths 340 nm and 440 nm, 380 nm and 620 nm, 320 nm and 530 nm, 460 nm and 570 nm, and 400 nm and 570 nm. As predicted rhabdomeric responses correspond to the log of the relative photon capture (Figure 2B-E), we mixed wavelengths in the following way to test for linearity:

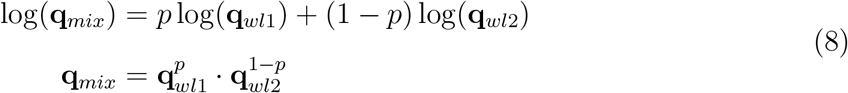

where **q**_*mix*_ is the calculated relative capture for the mixture of wavelengths, *p* is the proportion of wavelength *wl*1, **q**_*wl*1_ is the calculated capture of wavelength *wl*1, and **q**_*wl*2_ is the calculated capture of wavelength *wl*2. Using Equation 7, we fit the calculated captures for the mixture of wavelengths, as we did for the single wavelengths. For testing linearity of our responses, we used the mixtures at the luminant multiple of 1, as it provided good fits in our regression (see Figure S1D, J, P, S-W) and large calcium responses (see Figure S4 and Figure S5B-F).

For any analysis and modeling work, we used the calculated relative capture after fitting and not the target relative capture, we were aiming to simulate.

### Quantification of Imaging Data

All data analysis for *in vivo* calcium imaging was performed in Python using custom-made Python code and publicly available libraries. To correct our calcium movies for motion we performed rigid translations based on template alignment using the algorithm provided by the CaImAn package (Giovannucci et al., 2019). As a template for rigid motion correction, we used the average projection of the first ten seconds of every calcium movie during which we did not show any visual stimuli.

#### Image Segmentation

Region of interestes (ROIs) were selected automatically using a custom-made approach and verified manually. A standard deviation projection was taken of the complete image stack. We thresholded the projected image in three ways to identify pixels that are certainly part of a ROI, certainly part of the background, and possibly part of a ROI. These thresholded images were used to identify connected components (i.e. individual ROIs). Next, we applied a watershed transformation to obtain the individual ROIs. We discarded any ROIs of fewer than 5 pixels.

#### Signal Extraction

To extract calcium traces from our segmented images, we first took the average fluorescence of each ROI at each time point. We subtracted the mean background fluorescence - the mean fluorescence of all pixels that do not belong to any ROI - from each trace to remove background noise. To calculate the dF/F signal, we use as a baseline for our denoised traces the 5th percentile of a rolling 30 second time window. Finally, we smooth our dF/F signal with a Gaussian filter of size 0.32 seconds and a standard deviation of 0.08 seconds. We discarded ROIs, where the signal-to-noise (*SNR*) ratio was smaller than 1.5. The *SNR* was defined as the standard deviation of the signal during stimulation over the standard deviation of the signal before and after the start of stimulation (*SNR* = *σ*_*stim*_/*σ*_*baseline*_).

#### Peri-stimulus time histograms (PSTHs)

dF/F traces were aligned to the stimulus start times and averaged for each ROI. Amplitude measurements were taken on these averaged PSTHs for each ROI. As each stimulus was 0.5 seconds long amplitudes were calculated by taking the average dF/F response between 0.42 and 0.5 seconds after the stimulus onset and subtracting the average dF/F response 0.15 to 0.05 seconds before stimulus onset (i.e. the baseline). The max-dF/F signal of the spectral tuning curves was calculated by dividing the mean across all ROIs of the wavelength with the maximum response.

#### Signal sorting

In the case of the double NorpA rescues, we needed to sort our individual ROIs. To do this, we fitted the data to the log of the relative photon capture of the opsin each photoreceptor expresses. For example, in the case of the pR8 and yR8 NorpA rescues, we fitted each ROI to the log of the relative photon capture of Rh5 and Rh6 separately. Next, we assigned each ROI to the cell type according to which fit explained more of the variance. In our example, if the Rh5 fit is better than the Rh6 fit for a ROI, that ROI is a putative pR8 axon.

### Statistics

For all PSTHs and tuning curves, we show the empirically-bootstrapped 95% confidence interval to indicate significance. To obtain these intervals, we randomly resampled from our data 1000 times and recalculated the mean. Next, we took the 2.5% and 97.5% percentile of our 1000 samples.

We calculated the Bayesian Information Criterion (BIC) as follows:

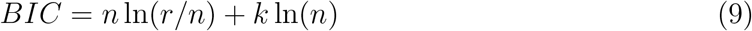

wheren *n* is the number of data points, *r* is the sum of the squares of residuals (deviations predicted from actual empirical values of data), and *k* is the number of parameters in the model.

### Data for Spectral Sensitivities

We obtained the spectral sensitivities for the opsins Rh3, Rh4, Rh5, and Rh6 from Salcedo et al. (1999). We fit the raw sensitivities to the following equation (Govardovskii et al., 2000):

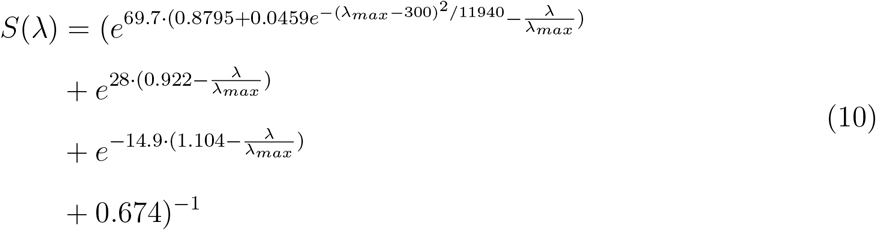

where *λ* is the wavelength, *λ*_*max*_ is the peak wavelength, and *S* is the fitted spectral sensitivity.

### Correlation Coefficients and Principal Components

To calculate the correlation coefficients of actual and predicted responses and obtain principal components of our spectral sensitivities, we first calculated the covariance matrix (**Σ**). For the spectral sensitivities, we calculated the covariance using a uniform Fourier frequency power spectrum as in Buchsbaum and Gottschalk (1983). For our predicted and actual responses, we calculated the covariance as follows:

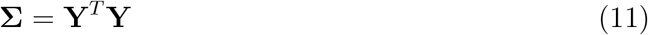

where **Y** are the responses of the different cell types across stimuli (*stimulus×cell−type*). We calculate the correlation coefficient matrix (**C**), as follows:

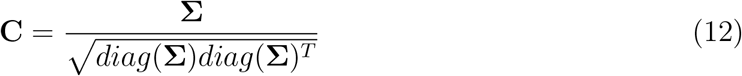

In order to decompose the opsin spectral sensitivities, we simply eigendecompose the covariance **Σ** to obtain eigenvalues and eigenvectors. This is equivalent to principal component analysis (PCA), where the eigenvectors correspond to the different components and the eigenvalues are proportional to the explained variance for each component. To construct principal component spectral sensitivity curves, we took the dot product of the opsins’ spectral sensitivities (*wavelength × opsin*) and the eigenvectors (*opsin × component*).

### Linear Regression

To assess chromatic tuning of our responses, we fit a linear regression model to our data:

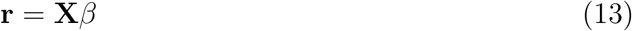

where **r** is the average amplitude response of a neuron type to the various stimuli in the flat background condition, **X** is the input space (i.e. the log(*q*) for each stimulus), and *β* are the associated weights for each input feature. Fitting was performed using 4-fold cross-validation. To improve numerical stability during the fitting procedure without biasing the end result, fitting was performed on a “whitened” input space (PCA whitening). After fitting, parameters were transformed back into “unwhitened” space. In order to assess goodness of fit for the different inputs, we calculated the 4-fold cross-validated *R*^2^ values for each input space.

Linear regressions that include intra-ommatidial R7/R8 opsin pairs together with at least one additional opsin type provide better fits than when exclusively considering intraommatidial R7/R8 opsin pairs as independent variables (Figure S5K-N). This is most obvious in the case of pR7 and pR8. These regressions are consistent with findings that both intra- and inter-ommatidial interactions shape opponent responses in R7/R8.

### Circuit Modeling

Given the hypothesized circuit architecture, we built a linear recurrent model described by the following equations:

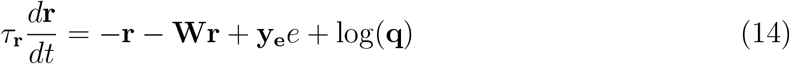

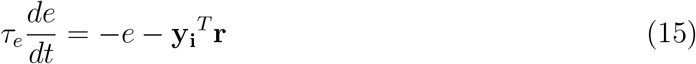

where **r** is a vector of the responses of the photoreceptor axons, **W** is the connectivity matrix for the direct inhibitory connections, *τ* is the time constant, **y**_**e**_ is a vector of the synaptic weight from Dm9 back to each photoreceptor, *e* is the response of Dm9, **q** is the relative photon capture, **y**_**i**_ is the synaptic weights from the photoreceptors to Dm9. All weights are positive, and the inhibitory or excitatory nature of the synapse is indicated by the sign.

We can simplify the above equation by setting 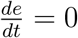, 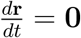 (i.e. steady-state condition), so that:

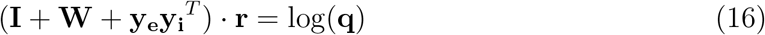

where **I** is the identity matrix. Using Equation 16, we fit the model to all our flat background data using least-squares and cross-validated our fits 4-fold.

**Table S2.** Synaptic count proportions for the different neurons in the recurrent circuit. Related to Figure 6.

| Input proportions to photoreceptors |  |  |  |
| --- | --- | --- | --- |
| Photoreceptor | direct input | Dm9 input |  |
| pR7 | 0.5 | 0.5 |  |
| yR7 | 0.5 | 0.5 |  |
| pR8 | 0.5 | 0.5 |  |
| yR8 | 0.5 | 0.5 |  |
| Input proportions to Dm9 |  |  |  |
| pR7 input | yR7 input | pR8 input | yR8 input |
| 0.21 | 0.14 | 0.32 | 0.33 |

The EM synaptic count proportions, we used, are in Table S2 (Reiser et al., 2019; Takemura et al., 2013). To normalize synaptic counts, we divided the synaptic count by the total number of synapses for each neuron. To change the gains of individual neurons using these fixed weights (synaptic count + gain model), we fit the Dm9 gain *c* and the photoreceptor gains **a** in the following equation using least squares:

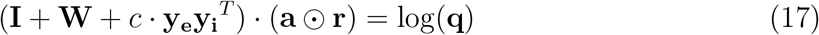

To compare synaptic gains to randomly drawn weights, we used the same Equation 17, and pulled random weights from a uniform distribution ranging from 0 to 1. Before fitting, the weights were normalized the same way the EM weights were normalized. We sampled a total of 10000 random weights to build a distribution of the cross-validated *R*^2^ values.

We used our synaptic count + gain model fits for our prediction of the spectral filtering curve and center-surround receptive field. The spectral filtering curve is the predicted response to individual narrow single wavelengths (instead of broader single wavelengths which we were able to test). The center-surround receptive field was normalized to each peak response. The center corresponds to the predicted response, when removing the Dm9 interneuron (*center* = (**I** + **W**) *⋅* (**a** ⊙ **r**)). The surround corresponds to the response to the input of each photoreceptor receives from Dm9 (*surround* = *c* ⋅ **y**_**e**_**y**_**i**_^*T*^ ⋅ (**a** ⊙ **r**)).

---

**Figure S1.**
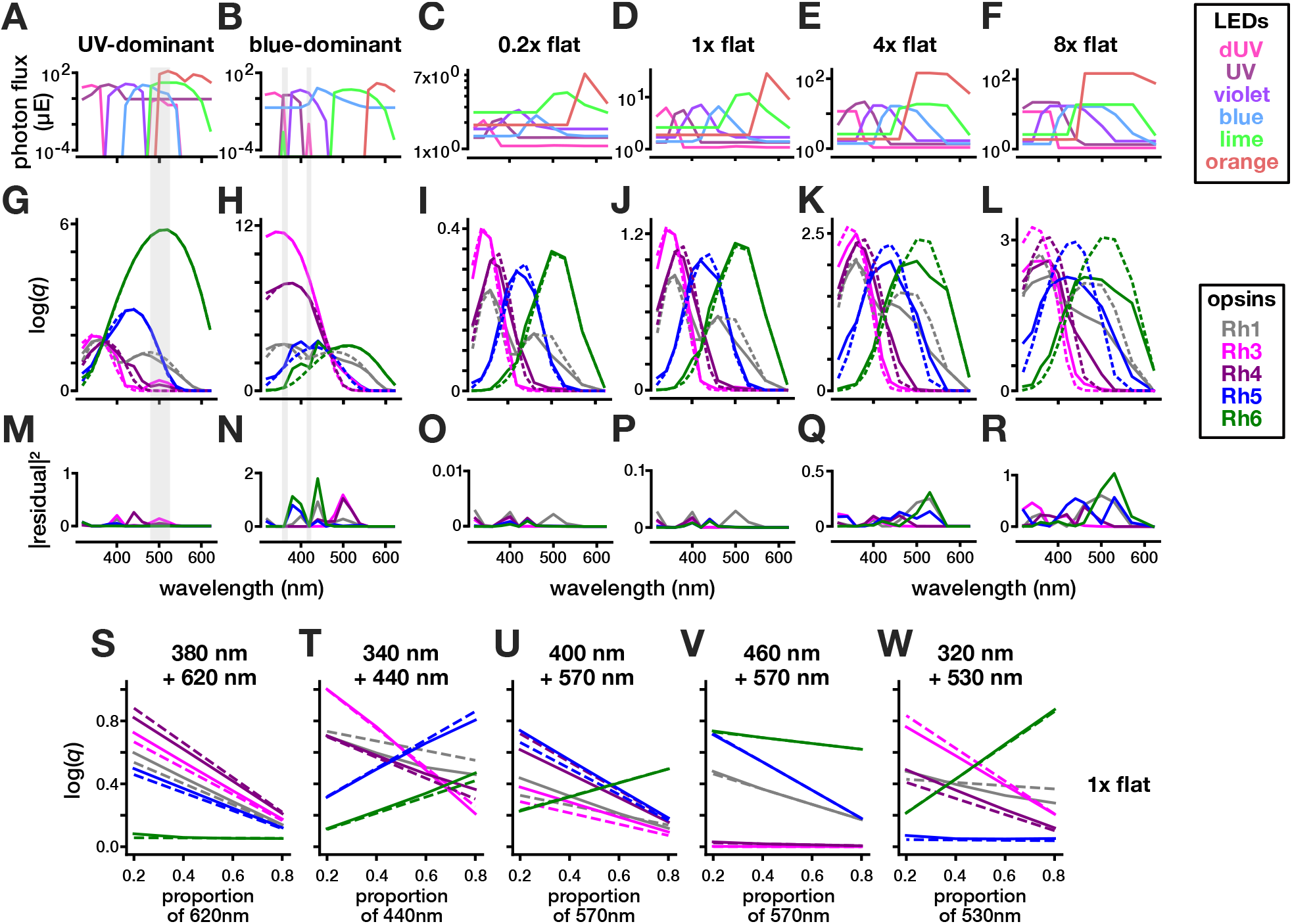
Contributions of individual LEDs to simulations of various wavelengths. Related to Figure 1. **A-F.** Relative photon flux of each of the 6 LEDs in the stimulation set-up used to simulate wavelengths across the spectrum for stimuli in the UV-dominant, blue-dominant, or flat background. **G-L.** Target log(*q*) of each opsin for a given simulated wavelength (dashed line) and log(*q*) of the best fit using our experimental setup (solid line) (see methods; Equation 7). **M-R.** The squared residuals calculated for target wavelengths and fitted wavelengths for all stimuli. The gray vertical shaded area in the UV- and blue-dominant background indicate wavelengths that were discarded when plotting spectral tuning curves. **S-W.** Target log(*q*) of each opsin for a given mixture of wavelengths (dashed line) and log(*q*) of the best fit using our experimental setup (solid line) (see methods; Equation 7 and Equation 8).

**Figure S2.**
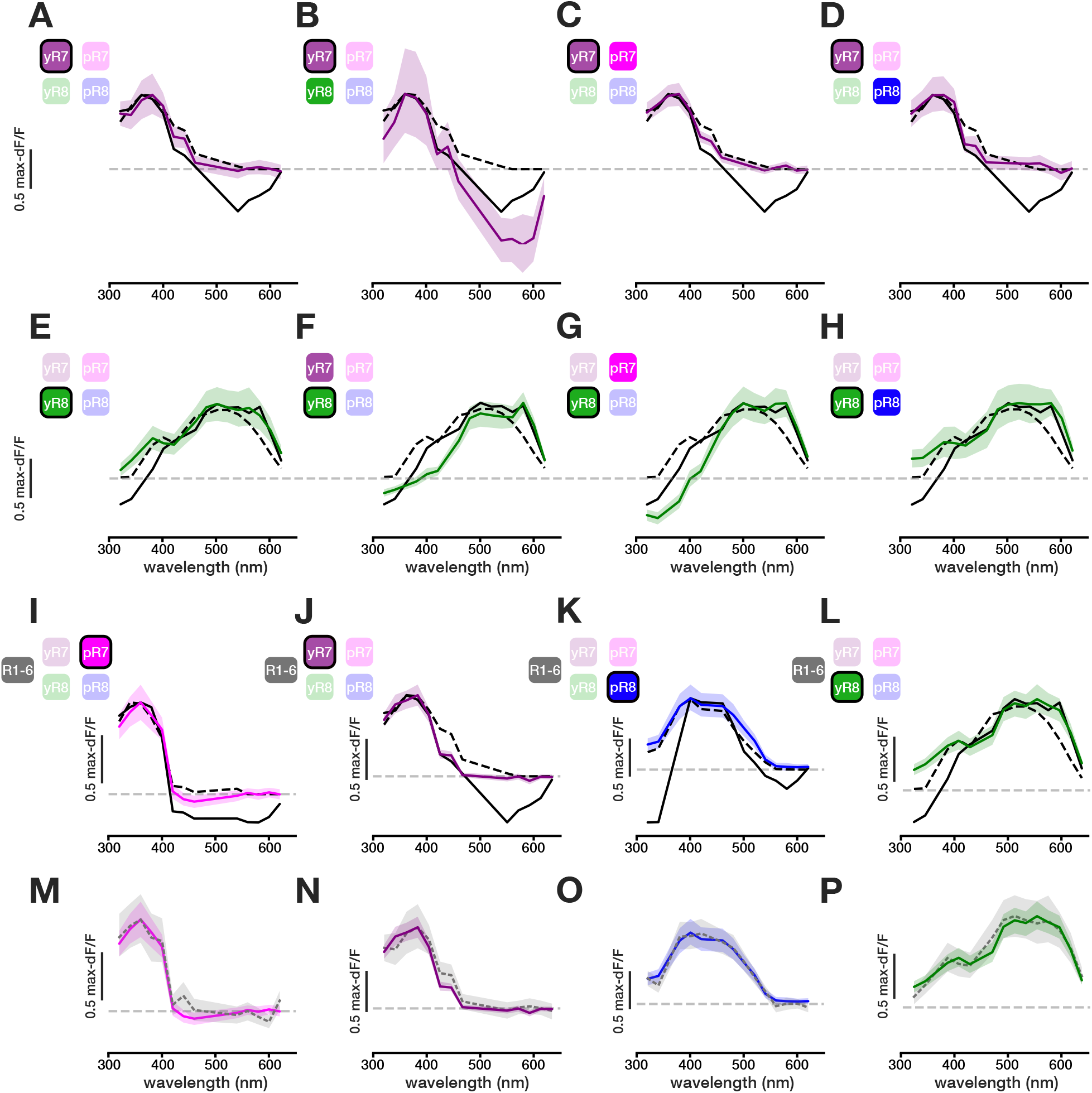
Pairwise NorpA rescues highlight sources of opponency in R7/R8. Related to Figure 3. **A-D.** Max-normalized responses of yR7 axons across wavelengths with NorpA restored in *norpa-* blind flies in yR7 alone, or in yR7 and the indicated photoreceptor type. (Ns= 132 ROIs (8 flies), 96(8), 135(12), and 51(5), respectively. **E-H.** Max-normalized responses of yR8 axons across wavelengths with NorpA restored in *norpa-* blind flies in yR8 alone, or in yR8 and the indicated photoreceptor type.(Ns= 126(6), 74(5), 48(6) and 26(4), respectively. **I-L.** Max-normalized responses of each photoreceptor axon type across wavelengths with NorpA restored in *norpa-* blind flies in select pairs of photoreceptor in the imaged photoreceptor and R1-6. (Ns= 150(6), 103(5), 85(7) and 95(4), respectively. **M-P.** Comparison of max-normalized spectral tuning curves of the single NorpA rescues and NorpA restored in the imaged photoreceptor and R1-6 showing no significant difference.

**Figure S3.**
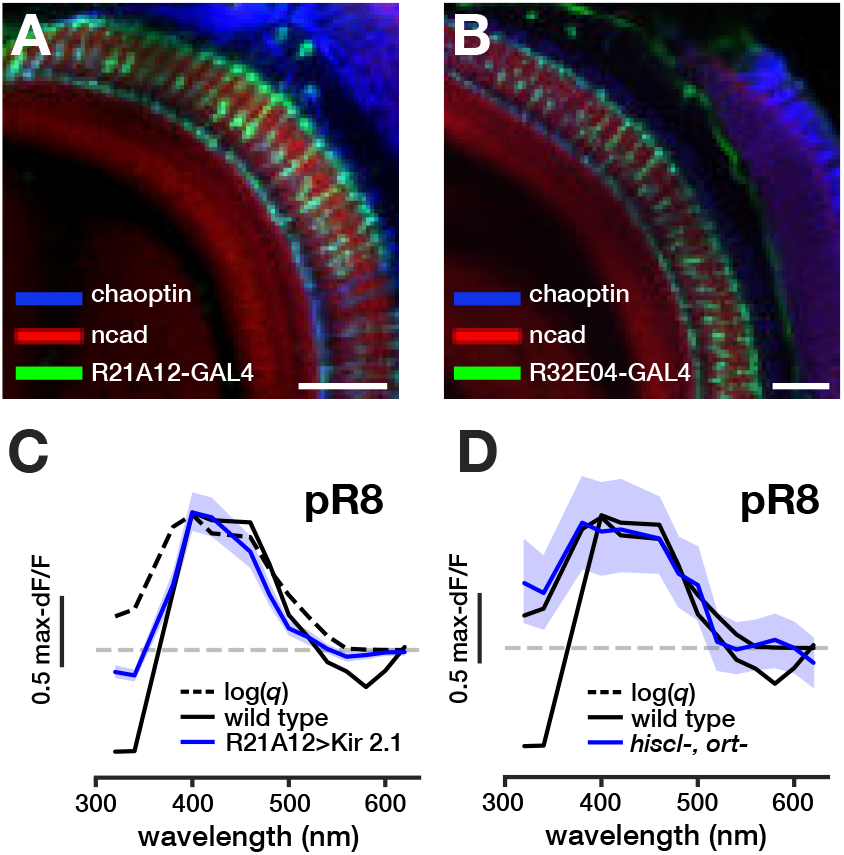
Immunolabeling of Dm9 and pR8 spectral tuning curves for various mutant background. Related to Figure 4. **A.** The optic lobe of the fruit fly stained for Dm9 (R21A12-Gal4, green), the neuropil (Ncad, red), and photoreceptor axons (Chaoptin, blue). Scale bar, 30 *µm*. **B.** The optic lobe of the fruit fly stained for Dm9 (R32E04-Gal4, green), the neuropil (Ncad, red), and photoreceptor axons (Chaoptin, blue). Scale bar, 30 *µm*. **C.** pR8 max-normalized spectral tuning curve when silencing Dm9 using the R21A12-Gal4 line (N= 158 ROIs (5 flies)).Dashed black lines represent log(*q*) and black lines represent the wildt type response. The shaded region represent the 95% confidence interval for the given spectral tuning curve. **D.** pR8 max-normalized spectral tuning curve in a *hiscl-*, *ort-* mutant (N= 153 (10)).

**Figure S4.**
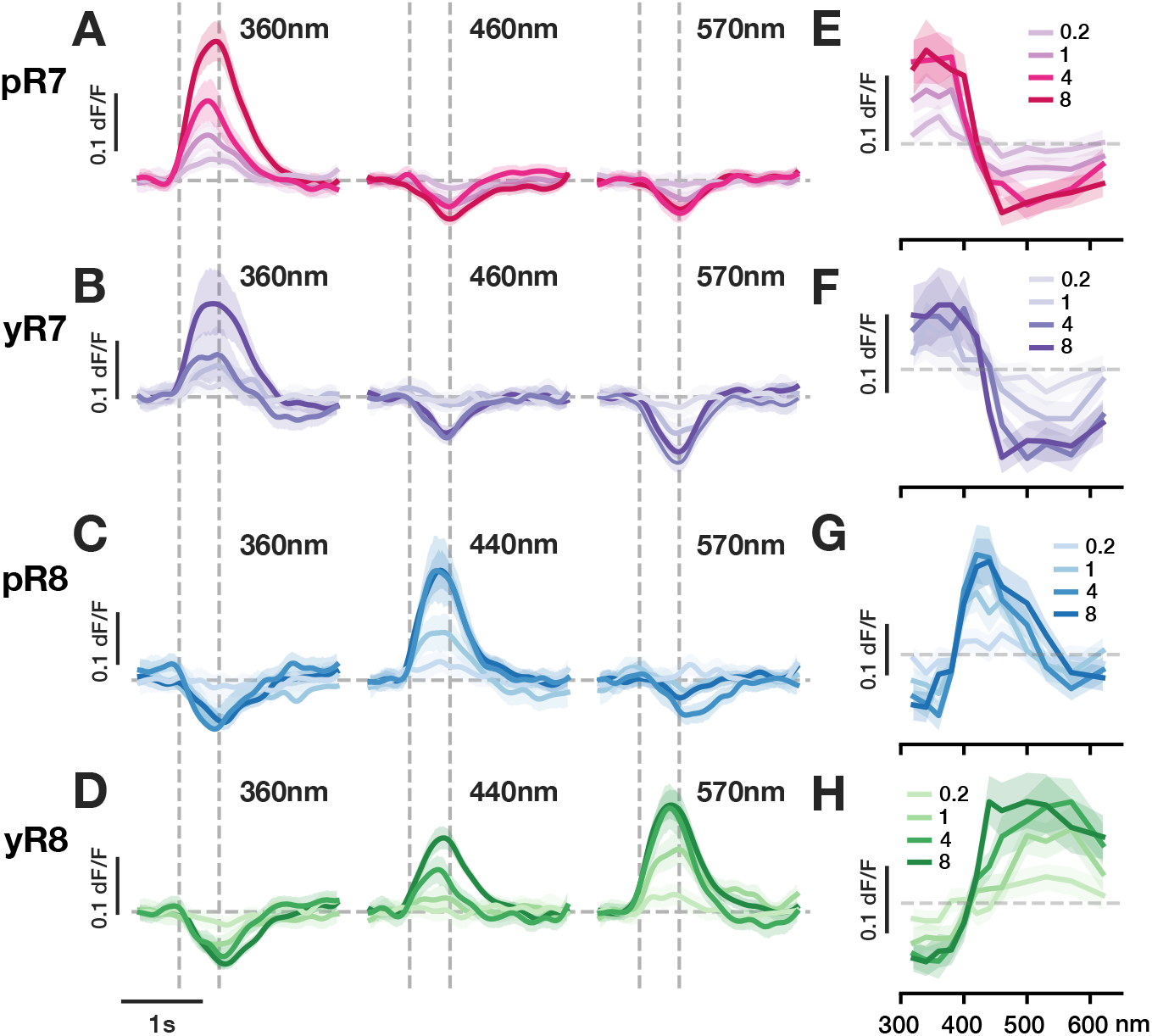
R7 and R8 spectral tuning to stimuli presented over a flat background spectrum. Related to Figure 5 and Figure 6. **A-D.** Average GCaMP6f responses of R7 and R8 axons in wild type flies to 0.5 second flashes of three simulated wavelengths over the flat background. Thick lines represent the mean, shaded region represents 95% confidence interval. Colored lines represent stimuli of four different luminant multiples. **E-H.** Tuning curves constructed using the amplitudes of measured responses of R7 and R8 axons from across the wavelength spectrum. Solid lines represent the mean, shaded region represents 95% confidence interval. Various colored lines represent stimuli of four different luminant multiples.

**Figure S5.**
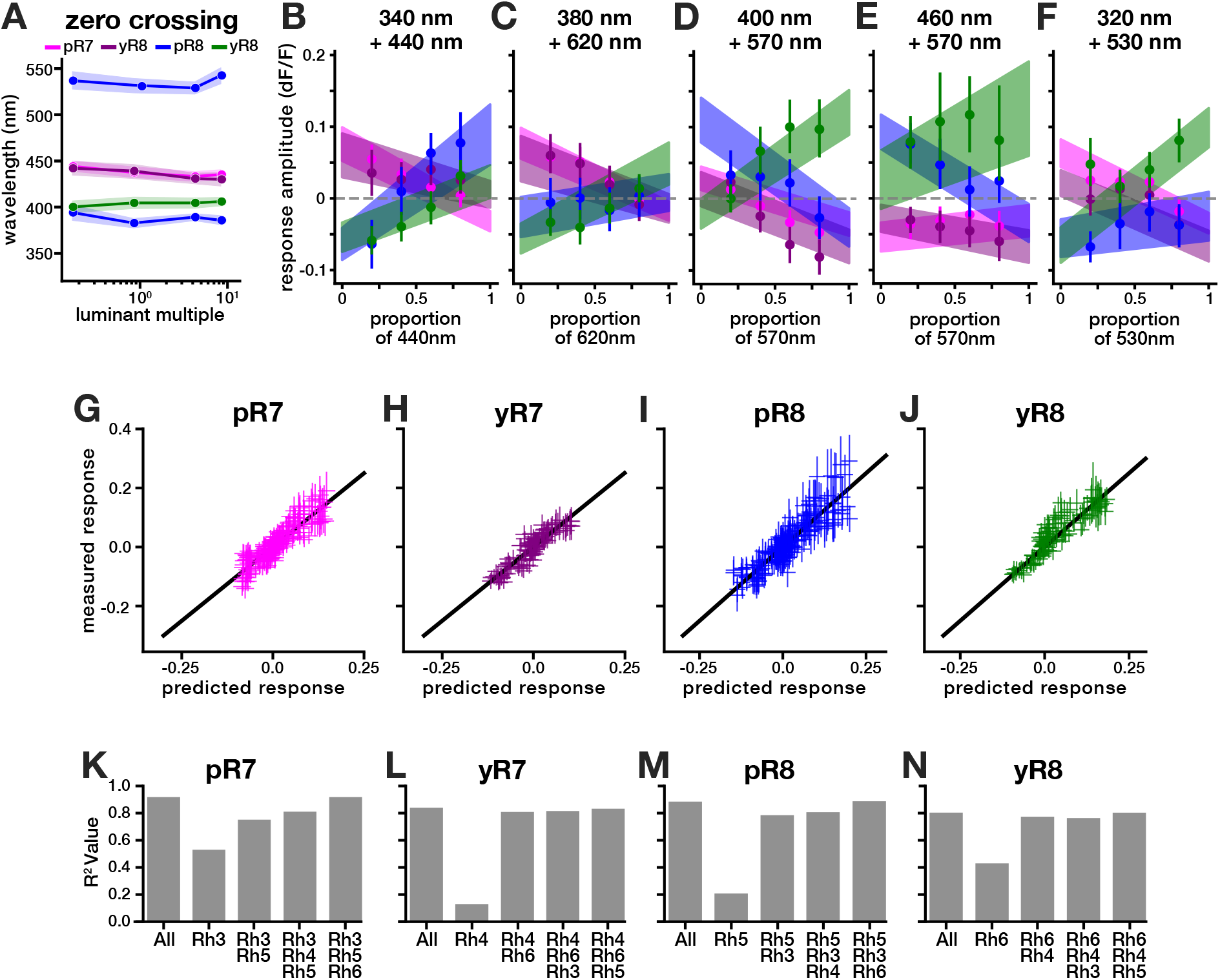
Measured responses in R7 and R8 axons in the 10 *µE* background result from linear combinations. Related to Figure 6. **A.** The estimated zero crossing points of the measured tuning curves in each R7/R8 photoreceptor. As pR8’s tuning curve has a trilobed form, it crosses the zero axis twice. **B-F.** Measured responses of R7 and R8 axons to mixed combinations of different wavelength stimuli (filled circles) compared to the linear prediction of responses to those stimuli (shaded region) (see methods; Equation 8). **G-J.** 4-fold cross-validated linear regression using the log of the relative photon capture of Rh3-6 and responses measured at the level of R7 and R8 outputs. **K-N.** *R*^2^ values when using varying opsin contributions in linear regressions to predict opponent responses in R7s and R8s.

